# Ancient Sheep Genomes reveal four Millennia of North European Short-Tailed Sheep in the Baltic Sea region

**DOI:** 10.1101/2023.06.26.544912

**Authors:** Martin NA Larsson, Pedro Morell Miranda, Li Pan, Kıvılcım Başak Vural, Damla Kaptan, André Elias Rodrigues Soares, Hanna Kivikero, Juha Kantanen, Mehmet Somel, Füsun Özer, Anna M Johansson, Jan Storå, Torsten Günther

## Abstract

Sheep are among the earliest domesticated livestock species, with a wide variety of breeds present today. However, it remains unclear how far back this breed diversity goes, with formal documentation only dating back a few centuries. North European short-tailed breeds are often assumed to be among the oldest domestic sheep populations, even thought to represent relicts of the earliest sheep expansions during the Neolithic period reaching Scandinavia less than 6000 years ago. This study sequenced the genomes (up to 11.6X) of five sheep remains from the Baltic islands of Gotland and Åland, dating from Late Neolithic (∼4100 calBP) to historical times (∼1600 CE). Our findings indicate that these ancient sheep largely possessed the genetic characteristics of modern North European short-tailed breeds, suggesting a substantial degree of long-term continuity of this breed type in the Baltic Sea region. Despite the wide temporal spread, population genetic analyses show high levels of affinity between the ancient genomes and they also exhibit higher genetic diversity when compared to modern breeds, implying a loss of diversity in recent centuries associated with breed formation. Finally, we see a potential signature of an even earlier, genetically different form of sheep in Scandinavia as these samples do not represent the first sheep in Northern Europe. Our results shed light on the development of breeds in Northern Europe specifically as well as the development of genetic diversity in sheep breeds, and their expansion from the domestication center in general.

## Introduction

Sheep (*Ovis aries*) were domesticated in the early Neolithic most likely somewhere in, or just to the east of, Anatolia (Zeder 2008). Sheep had spread into mainland Europe by around 8000 BP with human farming populations expanding from that region during the Neolithic period. Sheep reached Scandinavia with the same expansion, probably following the Danubian route, by around 6000 BP (Ryder 1984). The first Scandinavian farmers were associated with the Funnel Beaker culture (TRB; from German Trichterbecher) (Fischer 2002; Sjögren et al. 2019) and spread as far north as central Sweden. TRB was succeeded, starting around 5000 BP, by the Battle Axe Culture (BAC) which was a local variation on the pan-European Corded Ware Complex (CWC) (Malmer 2002). Studies of both CWC- and BAC-associated humans show that they have an ancestry component originating from the Pontic-Caspian steppe, associated with an expansion of the Yamnaya culture, which has not been observed in earlier individuals in Scandinavia (Allentoft et al. 2015; Haak et al. 2015; Malmström et al. 2019).

Today, northern Europe is home to the north European short-tailed sheep (NEST), a group consisting of 34 breeds (Dýrmundsson and Niżnikowski 2010). These sheep are characterized by a short and tapering tail, which is often also covered in wool. The group is considered to be more “archaic” than other European sheep (as reviewed by (Dýrmundsson and Niżnikowski 2010)). Some sources claim that this group is a relict of the first migrations of sheep into Northern Europe (Chessa et al. 2009; Rannamäe et al. 2020). Although microsatellite studies found variation between different breeds (Tapio et al. 2005) they show comparatively low levels of genetic diversity (Lv et al. 2022), which can be attributed to some NEST breeds’ long breed history and the fact that many breeds have historical census sizes of only a few individuals. NEST are now found in an area stretching from Iceland to Russia and are believed to have spread from Scandinavia through Norse Viking expansion (Ryder 1983; Dýrmundsson and Niżnikowski 2010). They are closely related to other Northwestern European breeds (Northern France, the British Isles, etc.), but most have small, isolated populations – both in mainland Scandinavia and Scotland and small islands like the Soay sheep of Saint Kilda (Dýrmundsson and Niżnikowski 2010; Stoffel et al. 2021). On the Baltic islands of Gotland and Åland there are three local sheep breeds (see Figure 1): The most common sheep on Gotland are of the breed “Gotland” (Swedish Board of Agriculture 2022) which was intentionally developed in the early 20th century as a commercial breed for their meat and pelts. Gute sheep, the traditional free-range breed on the island, went through a massive bottleneck in the mid-20th century as a result of the increase in popularity of the new Gotland sheep, but has recovered due to conservation efforts (Edberg 1986).

**Figure 1:**
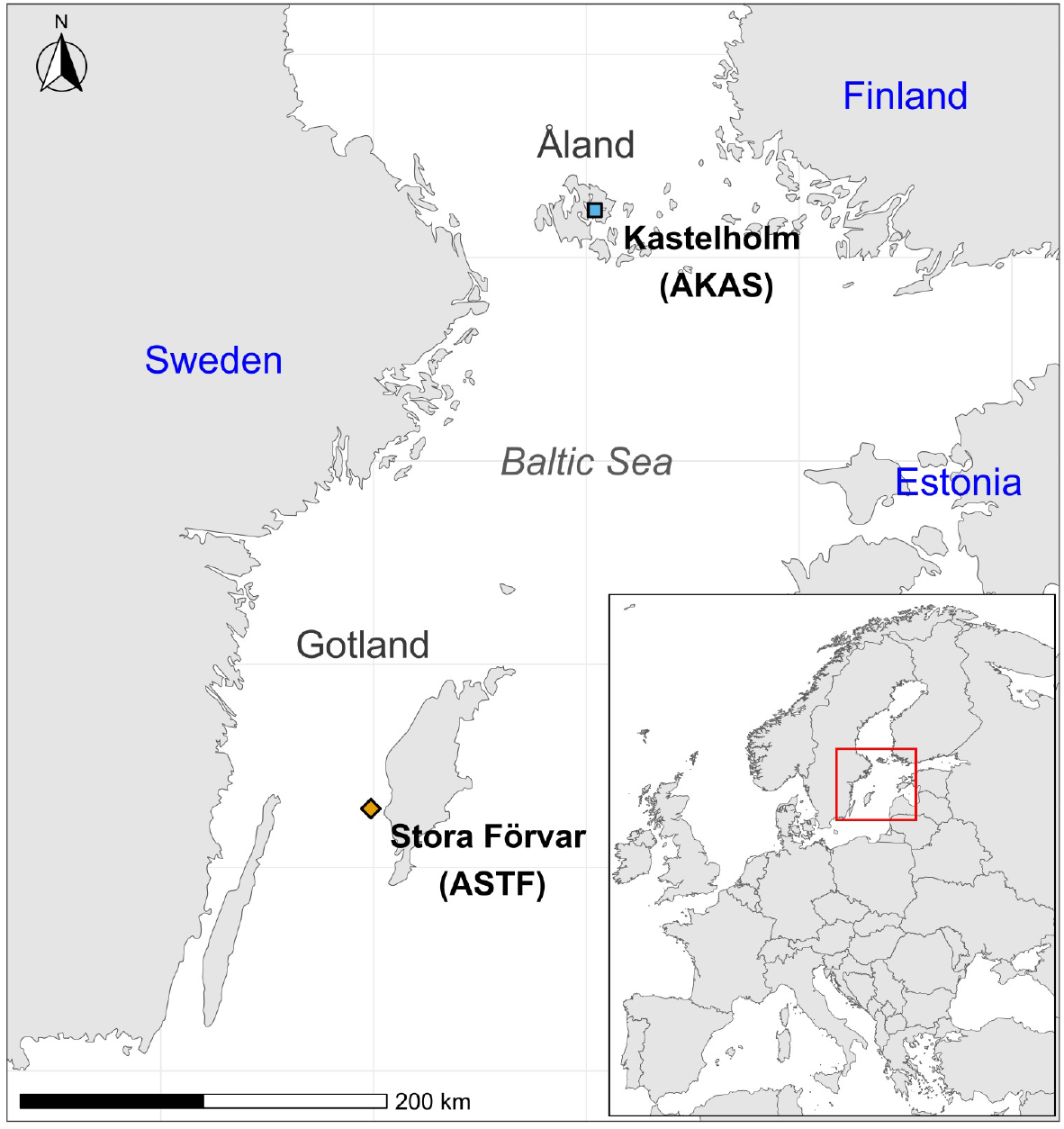
Map showing the Baltic Sea with some of the neighboring countries. Samples are marked by the blue square and the orange diamond. The inset map in the lower right corner shows the extent of the main map on a map of Europe.

Ancient DNA (aDNA) has developed into a powerful tool for studying the spread and history of domestic animals (Frantz et al. 2020). Archaeogenomic studies in sheep so far only include a few case studies in Anatolia (Yurtman et al. 2021), central Asia (Taylor et al. 2021), and Iran (Rossi et al. 2021). In northern Europe, aDNA studies have been restricted to uniparental markers (mitochondria and/or Y chromosomes) (Niemi et al. 2013; Rannamäe, Lõugas, Niemi, et al. 2016a; Rannamäe, Lõugas, Speller, et al. 2016b), which have limited power for understanding complex demographic processes due to their non-recombining nature (Mourier et al. 2012) and are also sensitive to sex-biased processes. Here, we generate ancient genomes of five Scandinavian sheep (up to 11.6X sequencing depth) dating to the Late Neolithic (on Gotland) and historical periods (on Åland). We investigate their relationship to modern NEST breeds and observe a surprising level of continuity over nearly 4000 years. However, a recent reduction in genetic diversity seems to coincide with the development of separate breeds in the last centuries as well as with the replacement of local breeds with larger meat breeds such as Texel and Suffolk by many farmers.

## Results

We successfully obtained genomic aDNA from five sheep bones excavated from islands in the Baltic Sea – three from the cave Stora Förvar on Stora Karlsö, an island near Gotland, and two from Kastelholm on Åland (Figure 1), locations with potential connections to both sides of the Baltic Sea. Two samples from Stora Förvar (ASTF hereafter) were radiocarbon dated to the Late Neolithic around 4000 cal BP while one sample from Kastelholm (AKAS hereafter) has been dated to historical times around the 16th century CE. The remaining samples that were not directly dated fit well within the same archaeological context as the other samples, so we assume they are of similar ages. DNA preservation ranged from 1.3% (AKAS001) to 47% (ASTF002) endogenous DNA allowing us to shotgun sequence the genomes up to 11.6X autosomal coverage (Table 1). The well preserved Late Neolithic material confirms Gotland and its limestone as an exceptional place for DNA preservation which has already facilitated several studies on human aDNA (Skoglund et al. 2014; Günther et al. 2018; Coutinho et al. 2020). The sequence data is showing fragmentation and deamination damage patterns as expected for aDNA (Supplementary Figure 1). Four individuals belonged to mitochondrial haplogroup B which is the most common haplogroup in European sheep while one individual (ASTF002) carried haplogroup A which is usually more common in modern Asian sheep breeds (Machová et al. 2022), but both haplogroups have been found in modern and ancient northern European sheep (Niemi et al. 2013; Rannamäe, Lõugas, Niemi, et al. 2016a; Rannamäe, Lõugas, Speller, et al. 2016b).

**Table 1:** Overview of samples sequenced for this study.

| Sample | Site | Autosomal Coverage | Sex | Age (2σ, 95.4 % probability) | Mitochondrial Haplotype |
| --- | --- | --- | --- | --- | --- |
| ASTF-001 | Stora Förvar, Gotland | 6.7153 | Female | 3957-3699 cal BP (Ua-71140) | B1a |
| ASTF-002 | Stora Förvar, Gotland | 11.6010 | Male | 4151-3936 cal BP (Ua-71139) | A1b |
| ASTF-003 | Stora Förvar, Gotland | 3.8368 | Male | Late Neolithic* | B1a2a1 |
| AKAS-001 | Kastelholm, Åland | 0.0703 | Female | 527-340 cal BP (Ua-71141) | B1a1 |
| AKAS-002 | Kastelholm, Åland | 1.3350 | Female | CE 1500-1550* | B1a2a |

BP=Before present (1950 CE)
\*Contextual dates

### Descriptive population genetic analysis

To place the prehistoric and historic sheep samples into the broader genetic diversity of sheep, we merged them with large genomic datasets of modern-day sheep breeds from Eurasia and Africa. We first started with a dataset derived from whole genome resequencing data (WGS dataset hereafter) with SNPs that were called with the involvement of outgroups (other species of the genus *Ovis*) (Supplementary Table 1) to reduce the potential impact of ascertainment bias towards commercial breeds. The full dataset included about 2.7 million SNPs called in 151 individuals from 31 Eurasian breeds as well as 10 Asiatic mouflons (*O. gmelini*) assumed to represent the wild ancestor of domestic sheep (Supplementary Table 2) (Naval-Sanchez et al. 2018; Deng et al. 2020; Li et al. 2020). We then projected the ancient sheep onto the major axes of variation (principal components, PCs) in the modern breeds. PC1 (explaining 6.31% of the variation) separates Asiatic mouflon from domestic sheep while further PCs sort out variation within domestic sheep (Supplementary Figure 2). PC2 (explaining 2.69% of the total variance) aligns breeds mostly along an East to West gradient while PC3 (2.32%) starts separating individual breeds forming a gradient between Solognote from France to Gotland from Sweden (Figure 2). In this analysis, the ancient samples fall with northern European sheep, close to NEST (Gotland, Finnsheep, OldNorwegian, and Shetland) but do not directly overlap with any of these modern breeds. This result is consistent with model-based clustering as implemented in ADMIXTURE (Alexander et al. 2009) where they are composed of similar ancestry components as modern NEST breeds but with some minor ancestry contribution from other groups including low levels of basal ancestry attributed to the same cluster as Asiatic mouflon (Supplementary Figure 5).

**Figure 2:**
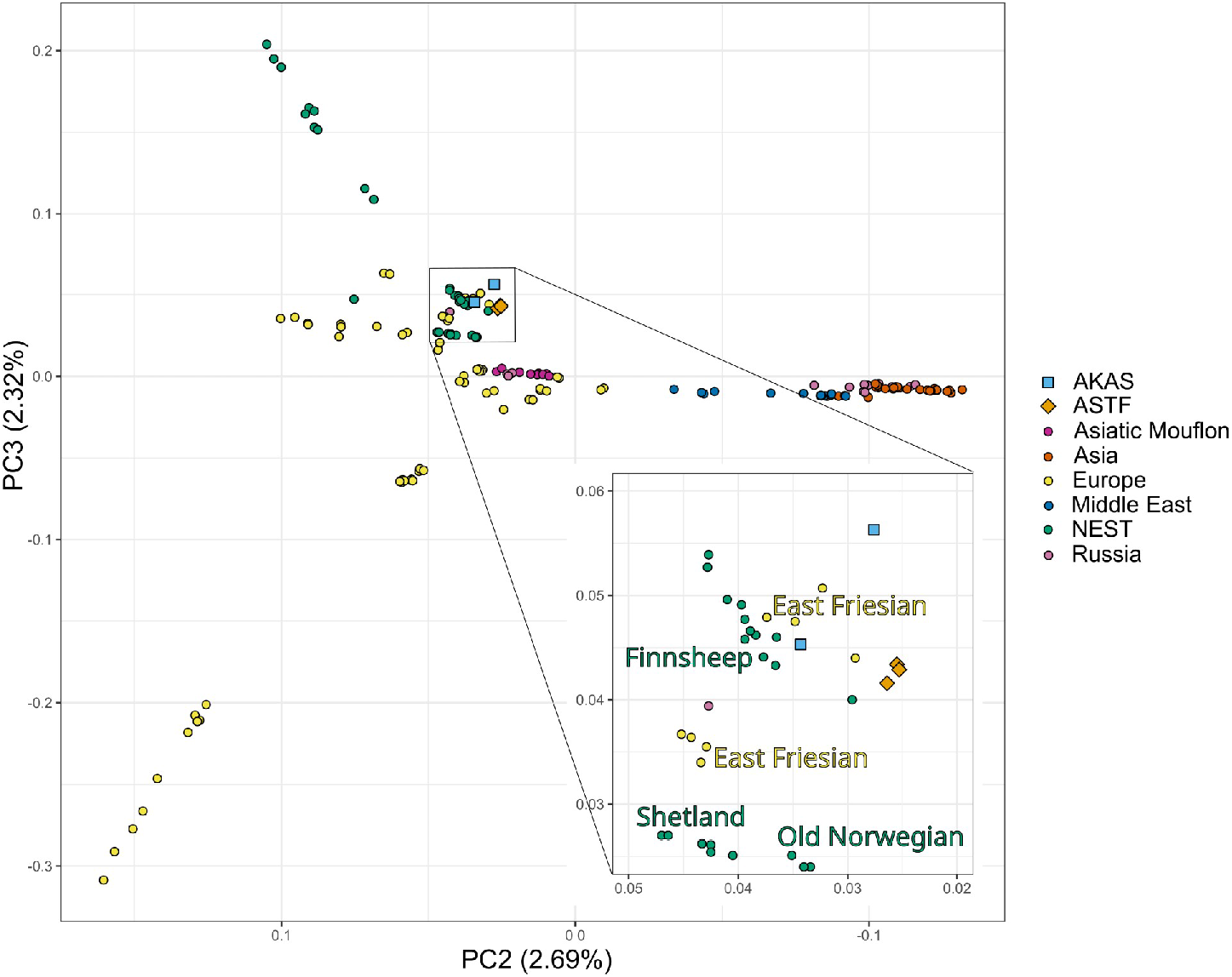
PCA biplot of the ancient Baltic sheep projected over the WGS dataset for PCs 2 and 3. PC2 matches the geographical distribution of modern sheep, while PC3 polarizes some of the NEST breeds from other European sheep. The inner square represents a zoom-in on the ancient samples.

### Comparison to a diverse set of NEST breeds

To further investigate the similarity of the ancient sheep to modern NEST breeds, we compiled a second dataset including more NEST breeds as well as breeds from Russia all genotyped on the Illumina Ovine Infinium® HD 600K chip (Rochus et al. 2018; Rochus et al. 2020a; Cao et al. 2021; Igoshin et al. 2022). The final dataset included 1010 modern individuals and 484,428 SNPs for analysis (SNPCHP dataset hereafter, Supplementary Table 3). Comparison with PCA and ADMIXTURE show similar results as for the WGS dataset, the ancient sheep display clear similarities to NEST breeds without directly matching any of them (Supplementary Figures 3, 4 and 6). This is particularly interesting as the SNPCHP dataset includes the two breeds from Gotland (Gotland and Gute) as well as the local sheep breed from Åland.

To identify the modern sheep breeds that share the most evolutionary history with the ancient sheep, we estimated shared drift using outgroup *f*_3_ statistics for both modern datasets. The estimates for ASTF and AKAS show a remarkably high correlation (Figure 3) despite being from separate islands and nearly 4000 years apart. For the WGS dataset (Figure 3A), we see the highest amounts of genetic drift shared with the NEST breeds (Finnsheep and Gotland showing the highest values) and a substantial gap to other breeds including other European breeds. For the SNPCHP data (Figure 3B), we see Åland sheep on top followed by Romanov and other NEST breeds. The inclusion of a more diverse set of NEST breeds as well as further breeds from Russia and Europe filled the gap between the top hits and all other breeds in this case. Finding Romanov sheep as one of the breeds with the highest amounts of shared drift with ancient Scandinavian sheep is remarkable as this Russian breed is assumed to have contributed to the gene pool of modern NEST breeds (Ghoreishifar et al. 2021). These outgroup *f*_3_ results can be corroborated by a maximum likelihood tree based on the allele frequencies (Pickrell and Pritchard 2012; Molloy et al. 2021) in the different breeds (Figure 4). In the OrientAGraph results, ASTF and AKAS group together with Åland as the closest group followed by Romanov who cluster with NEST breeds instead of with other Russian breeds.

**Figure 3:**
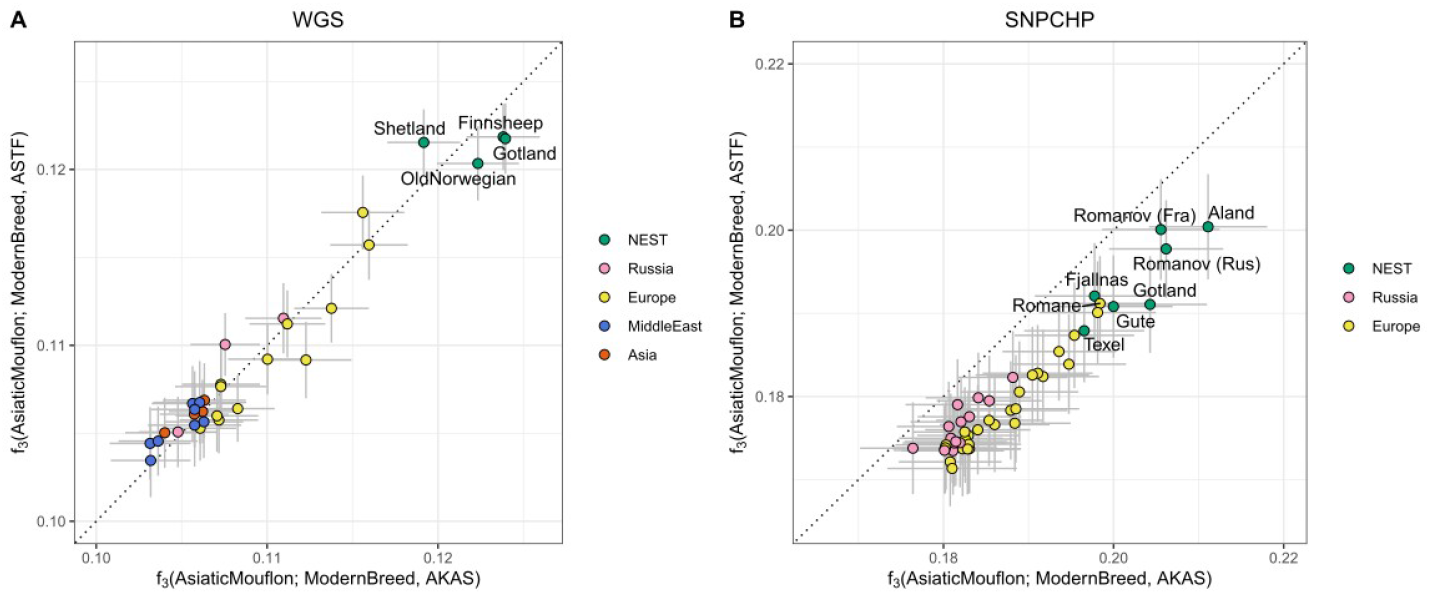
Outgroup *f*_3_ biplot measuring the shared drift of the ancient Baltic sheep from both sites/time periods versus modern breeds. Error bars show two block-jackknife standard errors. (A) WGS dataset, (B) SNPCHP dataset.

**Figure 4:**
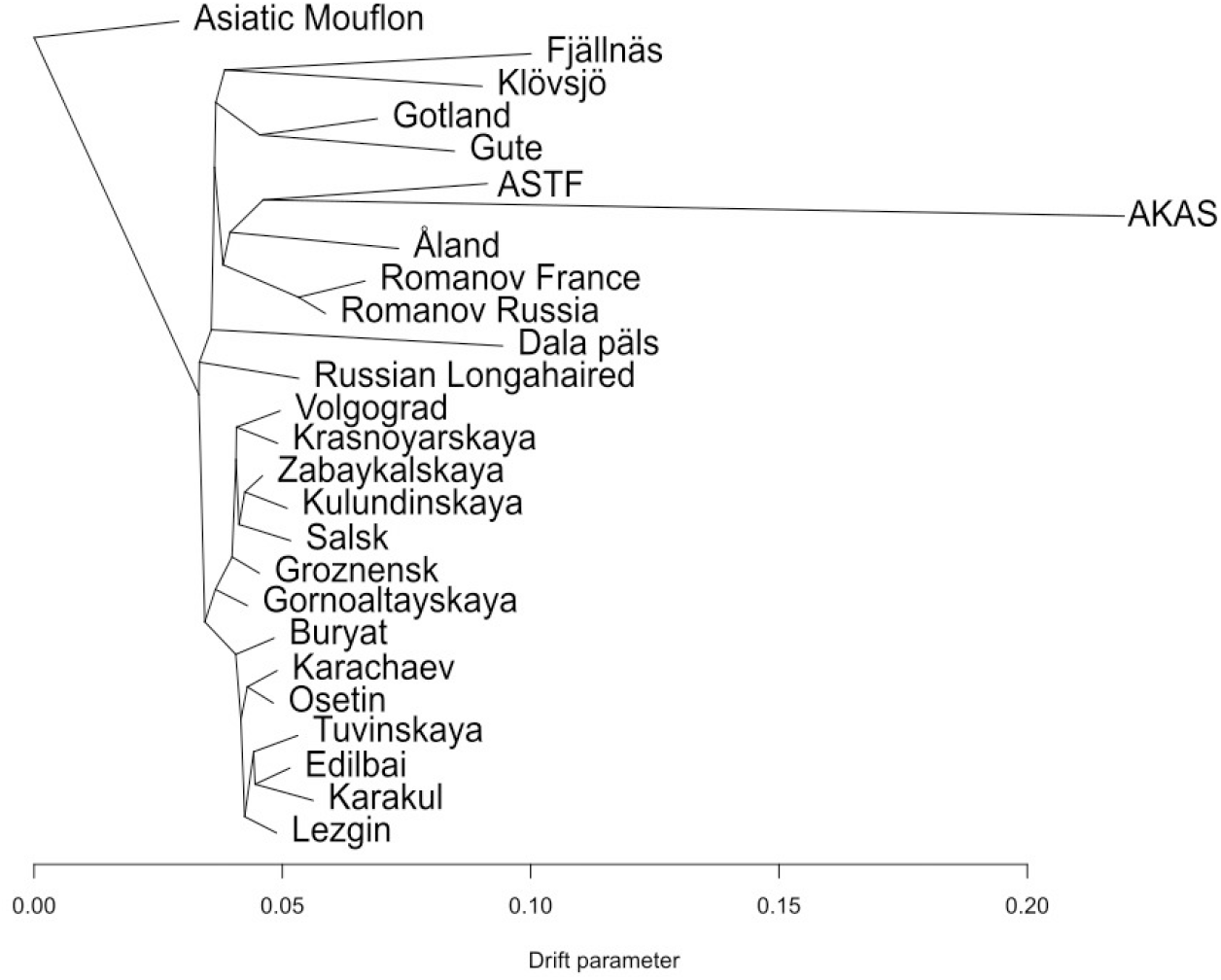
Maximum Likelihood Tree (using OrientAGraph (Molloy et al. 2021)) based on the covariances of allele frequencies in the SNPCHP data restricted to NEST and Russian breeds. The tree was estimated with OrientAGraph not allowing any migration edges.

While these results identify certain NEST breeds as closest modern-day relatives to ASTF and AKAS, they cannot be used as evidence for them representing the direct ancestor of any modern breed. An explicit test for population continuity (Schraiber 2018) rejected continuity for all breeds included in both datasets (Supplementary Tables 4 and 5). It needs to be noted that this test is considered extremely strict and even minor levels of independent evolution or gene flow lead to a rejection of the continuity model (Schraiber 2018). In agreement with the OrientAGraph results (Figure 4), models of discontinuity between modern and ancient samples estimate a substantial degree of private drift in the ancient populations which might be attributed to a combined effect of them representing island populations and aDNA data quality. Among all breeds, we also see that Romanov, Åland sheep and other NEST breeds show the lowest levels of divergence, consistent with the results of the outgroup *f*_3_ and OrientAGraph analyses (Supplementary Tables 4 and 5).

### Development of genetic diversity over time

The good preservation of the sheep remains sequenced in this study allows us to estimate genetic diversity for the prehistoric and historic populations and to compare them to the same estimates in modern breeds. As aDNA damages and other properties of the data complicate calling diploid genotypes, we chose an approach where we count pairwise differences between individuals from the same group at a set of biallelic sites known to be polymorphic (Skoglund et al. 2014). The result should provide an estimate correlated with the expected heterozygosity in each population and as the data is restricted to transversions, the results should not be driven by post-mortem deamination damages. The pairwise mismatch results for modern populations are largely similar to what has been estimated from whole genome resequencing data (Lv et al. 2022) with Asiatic mouflon displaying the highest diversity of all sheep populations, consistent with a domestication bottleneck, and more variation between particular breeds than between continental groups (Figure 5). Notably, both ASTF and AKAS show relatively high levels of diversity. They are significantly higher than all estimates for NEST and Russian breeds in the WGS dataset despite seemingly belonging to the same gene pool as modern NEST breeds. This suggests a very recent loss of genetic diversity in Northern European sheep in the last 500 years. The comparison between modern NEST breeds and ASTF/AKAS is similar when using the SNPCHP dataset although many other European and Russian breeds show higher values in this case (Supplementary Figure 8). Even if this broader pattern may be driven by ascertainment bias in the SNP array data, observing consistent results in both datasets suggests the robustness of the comparison between ASTF/AKAS and NEST breeds and the inference on recent diversity loss. The relatively high coverage data (11.6X) for ASTF002 also allowed us to call diploid genotypes (Prüfer 2018) and to estimate runs of homozygosity for this individual and compare the results to modern sheep breeds. The general pattern of ROH is comparable to the results for the pairwise mismatches with ASTF002 falling towards the lower end of modern NEST groups but we do not want to draw broader conclusions from this single sample (Supplementary Figure 9).

**Figure 5:**
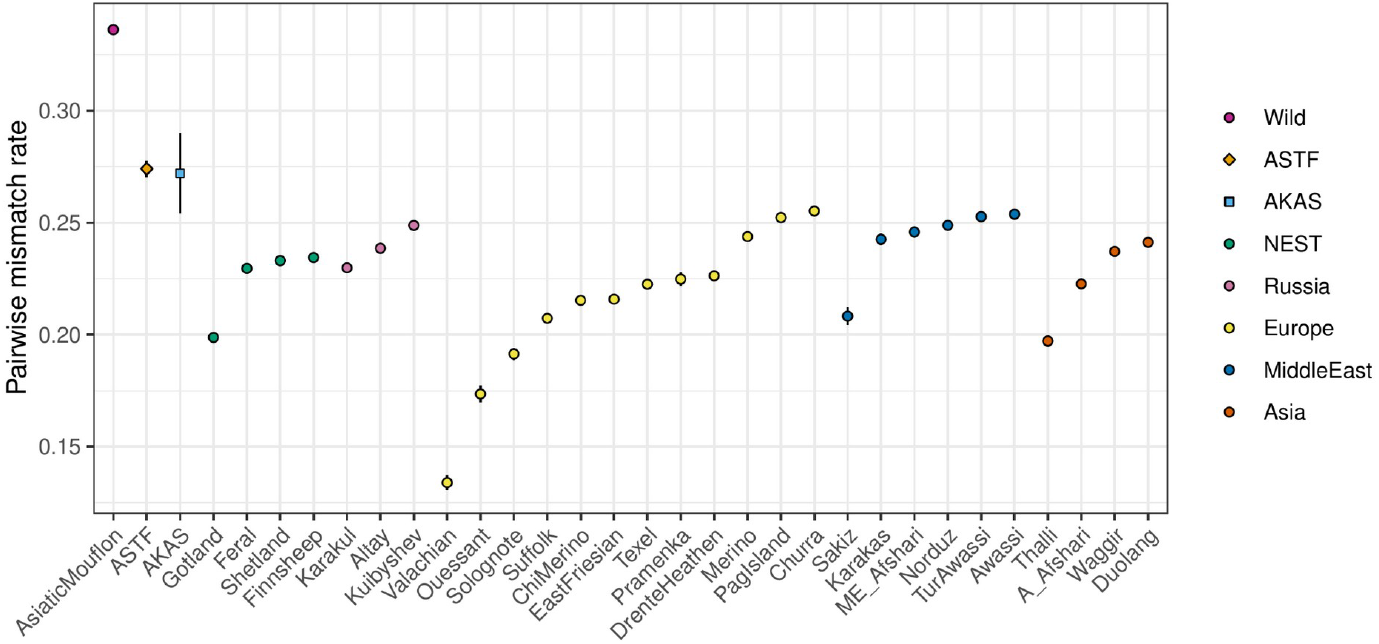
Pairwise mismatch rate between randomly sampled individuals from the same group. This analysis was performed on the WGS dataset. Error bars indicate two block-jackknife standard errors (for most groups, the error bars disappear behind the symbols). This analysis is based on transversions only to avoid post-mortem damage driving the differences between ancient and modern populations.

### Reconstruction of admixture graphs

To display the complex relationship of the ancient sheep with NEST and Russian breeds, we systematically explored the space of possible admixture graphs with admixtools2 (Maier et al. 2023). Admixture graphs are an effective way for displaying the relationships between populations and correlations between their allele frequencies. Despite their illustrative value, they should not be interpreted as demographic models displaying a full picture of the past. The most informative admixture graph corroborates some of the results found in other analyses (Figure 6). Pairwise comparisons between the best models for 0 to 5 admixture events (Supplementary Figures 10a-e) reveals that the fit of the models does not improve substantially (p<0.01 in pairwise comparisons to the next more complex model) beyond 3 admixture events (Supplementary Table 7). The European mouflon splits first as a representative of the Mediterranean lineage which is followed by Russian and NEST breeds splitting off. ASTF and AKAS are modeled as admixed sister groups with 84% of their ancestry coming from a source related to Åland sheep and 16% from an unobserved population basal to all other domestic sheep. This is consistent across all models including admixture events. For more than 3 events, the unsampled source turns from a basal group to a sister branch of all other sheep (Supplementary Figures 10d-f). The ADMIXTURE results also indicated low levels of basal ancestry in ASTF and AKAS (Supplementary Figure 5). The two other admixture events model the Asian Karakul as a mixture between a population related to ASTF/AKAS (68%) and Russian Longhair (32%) while Romanov are modeled as admixed between a population related to Karakul (24%) and Åland (76%).

**Figure 6:**
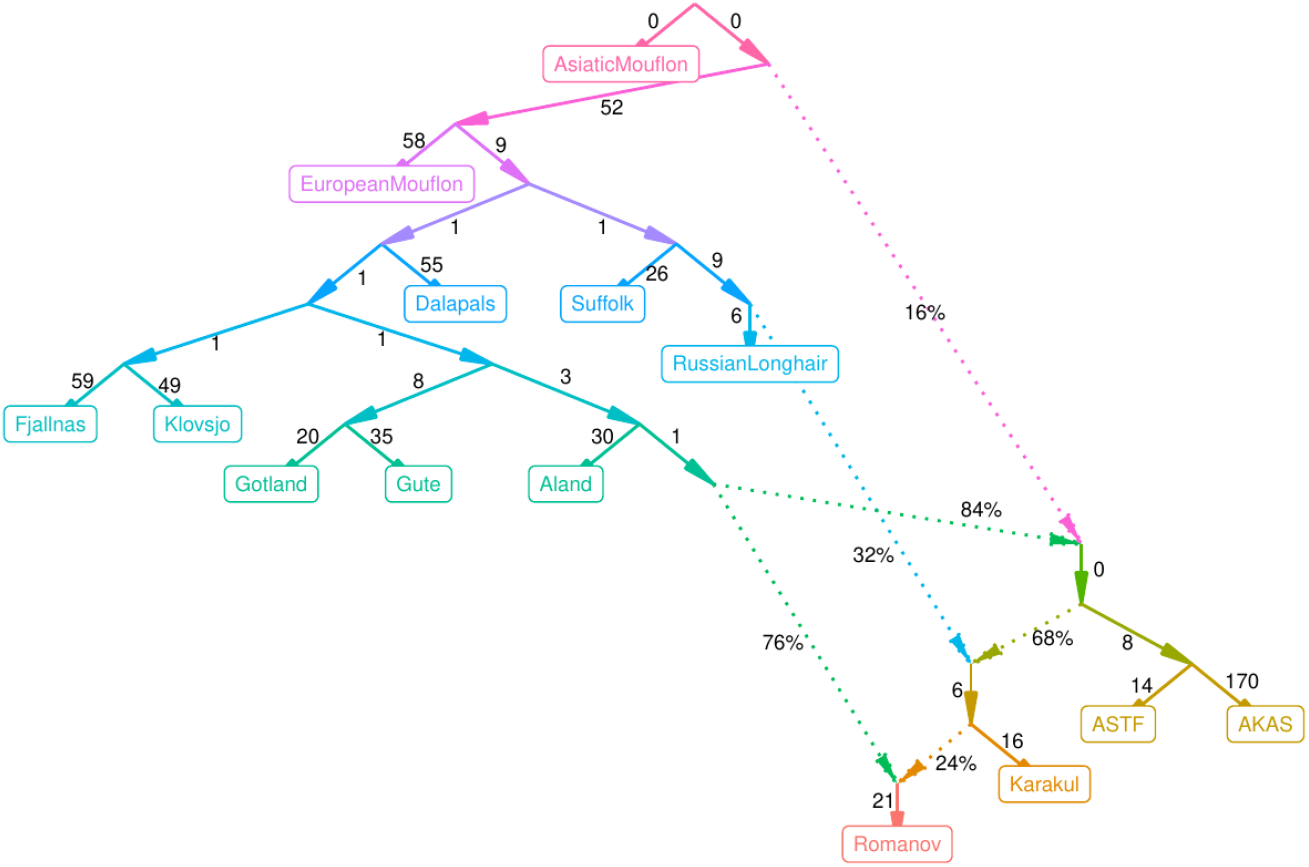
Well fitting admixture graph summarizing the relationship between selected NEST and Russian breeds based on correlations between their allele frequencies. The graph was constructed with three admixture events using the SNPCHP dataset. Graphs with other numbers of gene flow events are in Supplementary Figures 10. Comparison of the fit between models can be found in Supplementary Table 7.

It appears counterintuitive that the ancient ASTF/AKAS are modeled as an admixed group between an unsampled group and a modern breed even though this seems to be a consistently good fit to the data. Therefore, we performed a second search with a slightly reduced dataset restricting the search space by requesting that ASTF and EUR are not admixed as representatives of the Northern and Western European expansion of sheep. In this case, the model fit does not improve beyond one admixture event (Supplementary Table 8). The resulting model treats ASTF as the first population that splits off all other groups followed by European mouflon (Supplementary Figure 11b). Instead, Karakul are modeled as a group admixed between a population related to Russian Longhair (89%) and a group basal to all domestic sheep (11%). In the unconstrained model Karakul had also received some level of basal ancestry through ASTF/AKAS (Figure 6), representing a consistent signal of basal ancestry in some breeds. Higher numbers of admixture events result in qualitatively similar results as for the unconstrained model (Supplementary Figures 11a-f).

### The wool phenotype in the ancient sheep samples

To investigate the fiber quality in the ancient sheep, we checked two loci that were associated with wool fiber thickness or quality according to the literature (Demars et al. 2017; Lv et al. 2022). First, we genotyped the insertion of an antisense *EIF2S2* retrogene into the 3’ untranslated region (UTR) of the gene *IRF2BP2* by mapping to modified versions of the Oar3.1 reference genome with and without the insertion (Demars et al. 2017; Rossi et al. 2021). As multiple copies of the retrogene exist in the genome, we can only use reads spanning the insertion breakpoints as diagnostic. Using the reference genome including the insertion, only a single read for all five individuals maps across an insertion breakpoint, by only 3 bp out of which two are mismatches to the reference sequence (Supplementary Figures 12). In the modified reference without the insertion, two reads are crossing the breakpoint (Supplementary Figures 12). While the number of reads is small in both cases, we do not consider this strong evidence for presence of the derived allele. The ancestral state was also found at the SNP rs161553028 (Supplementary Table 9), also located in the 3’ UTR of *IRF2BP2* and correlated with fiber thickness according to (Lv et al. 2022). Finding the ancestral state for both mutations is not surprising as most investigated modern NEST individuals are also ancestral at these mutations (Lv et al. 2022). While these results suggest some coarser wool in the ancient sheep, we cannot make a statement whether they were used as a source of fiber in addition to meat and potentially milk.

## Discussion

Despite only representing five individuals spread across almost four millennia, the insights gained from our aDNA data highlight the power of studying the recombining part of the genome in temporal data for our understanding of demographic processes. The individuals from the two Baltic islands of Åland and Gotland show remarkable genetic similarities to each other suggesting that they were part of the same gene pool that has been present in the Baltic Sea region for the last four millennia. A long continuity of sheep in the Baltic Sea region since at least the Late Bronze Age had already been suggested based on uniparental data (Niemi et al. 2013; Rannamäe, Lõugas, Speller, et al. 2016b). Our study now extends this observation by another millennium on the autosomal level.

We see the highest level of similarity between our ancient sheep, including the Late Neolithic, and the modern Åland breed. While the ancient genomes display a high similarity to NEST breeds in general, none of the modern breeds is a direct match for a population with uninterrupted continuity. This suggests that other genetic processes and input from outside of Scandinavia as well as potentially unsampled Northern European populations were involved in the development of these separate breeds during the last few centuries. In the 16th century these sheep were just old traditional types of sheep in Scandinavia, but now most sheep used for production are crosses between fine-wool and heavier meat sheep, and some level of cross-breeding with other breeds has been documented for almost all NEST breeds (Dýrmundsson and Niżnikowski 2010). The drop in genetic diversity after the AKAS individuals lived, i.e. since the 16-17th century, can probably be attributed to the development of separate breeds involving more controlled mating among individuals and/or to the reduction in the number of local sheep during the last centuries when larger meat-type breeds became more common. Simultaneously, it is remarkable that the diversity stayed constant between ∼4000 calBP and the 16th century CE suggesting that the mating patterns have not changed substantially during this time period. This is also in contrast to domestic plants such as sorghum for which a continuous reduction of genetic diversity seems to have taken place over the last millennia (Smith et al. 2019). A recent drop in genetic diversity of Northern European sheep has also been found in Bayesian Skyline plots based on uniparental markers although these suggested a lower level of diversity before the Medieval period (Niemi et al. 2018).

Some of our results can be interpreted as indication for an even earlier expansion of genetically distinct sheep that predated the Late Neolithic individuals from ASTF. ADMIXTURE (Supplementary Figure 5), OrientAGraph (Supplementary Figure 7) and qpGraph (Figure 6) all suggest small levels (<20%) of basal ancestry in the ancient sheep genomes that cannot be represented by any of the sampled groups. This would suggest that such ancestry has been diluted by later processes. The consistency of the pattern across different datasets and analyses yields some credibility to this hypothesis. The Late Neolithic sheep postdate the arrival of farming practices and sheep in Scandinavia by more than 1000 years, which would represent the Funnel Beaker culture (TRB). The Late Neolithic sheep also postdate the Battle Axe Culture (BAC), a Scandinavian variant of the Corded Ware horizon which has been shown to introduce a different type of genetic ancestry from the Pontic-Caspian steppe into human populations (Malmström et al. 2019). The Late Neolithic and Early Bronze Age were turbulent periods showing large transformations in the human gene pool all over Europe (Allentoft et al. 2015; Haak et al. 2015). Similar processes may have taken place in human dependent species around this time. In horses, the expansion of the so-called DOM2-lineage dates to this time (Fages et al. 2019; Librado et al. 2021) and it is possible that sheep populations have also seen secondary expansions, potentially originating in the Eurasian Steppe and associated with the secondary products revolution and the expansion of wool economies around this time (Sherratt 1983; Chessa et al. 2009; Marciniak 2011). Our current dataset, however, does not allow us to directly test this hypothesis as we lack both early or middle Neolithic Scandinavian sheep genomes as well as a description of a potential source gene pool for this secondary expansion. Future studies combining genomic and other bioarchaeological data across time and space will offer insights into these questions and processes for sheep in Scandinavia and elsewhere as well as for other domestic species.

## Materials and Methods

### Site description and archaeological samples

Stora Karlsö is a small island with several caves located approximately seven kilometers west of the island of Gotland, Sweden. The biggest cave on Stora Karlsö is called Stora Förvar and shows some of the earliest signs of human habitation on Gotland. The cave site Stora Förvar was excavated between 1888 and 1893 (Schnittger and Rydh 1940). Around 9300 years ago (cal BP), hunter-gatherers, mainly subsisting on marine hunting and fishing, lived in the cave (Lindqvist and Possnert 1997; Lindqvist and Possnert 1999). The cave was in repeated use except for a hiatus between 7600 cal BP and 6000 cal BP when the cave appears to have been abandoned. After ∼6000 cal BP the cave again was used and some of the oldest dates of domesticated animals have been recorded from the cave around 6000 cal BP (Lindqvist and Possnert 1997; Rundkvist et al. 2004). After the Neolithic period the cave exhibits a continuous use, but with varying intensity, until the Iron Age and historical times (Schnittger and Rydh 1940). The Stora Förvar cave is approximately 25 m deep and at the time of the excavations the cave was filled with “cultural layers” with a thickness of up to 4 m (Schnittger and Rydh 1940). The cave was excavated in sections (Sv. Parceller, labeled A-I, from the entrance and to the inner part of the cave) and in 0.3 m spits, i.e. the equivalent of a Swedish “foot” (Schnittger and Rydh 1940). Three right radii of sheep from Stora Förvar were included in the study (Supplementary Table 6). The samples were recovered in section I which was in the innermost parts of the cave where seven layers were noted. Thus, the thickness of the sediments was approximately 2.1 m in section I. The section was the last one to be excavated in 1893. The samples originate from layers 4 and 5 which were the deepest layers exhibiting bones from domesticated animals. Layer 6-7 in section I exhibits Mesolithic finds but with some recent intrusions of finds, e.g. pottery sherds. Two radiocarbon dates in section I fall in the Mesolithic (Apel et al. 2018). One sample was recovered from the cave wall c. 53 cm above the (excavated) cave floor, i.e. approximately in layer 6. Two sheep bones in section I, also recovered in the cave wall (c. 1.7 and 2 m from the floor) were previously radiocarbon dated to the Early Iron Age, roughly to the second century AD (Apel et al. 2018). So far, the oldest dating of sheep at Stora Förvar is ∼5600-6000 cal BP (Rundkvist et al. 2004).

Kastelholm castle is located on the main island of the Åland archipelago, a self-governing Swedish-speaking county of Finland. The castle was built at the end of the 14th century. The oldest written record is from 1388 and the castle became the administrative center for the Åland Islands. The castle burnt down several times in both the 16th and 17th centuries but was rebuilt and expanded. The landscape was modified and especially the area at the shore closest to the castle. The castle lost its administrative functions in the 1630’s and by 1747 the castle had been abandoned and became a ruin (Palamarz 2004; Kivikero 2020). Over the years many noblemen and their bailiffs held the castle, both Swedish and Danish. Sheep meat, wool and other products were collected as tax from the archipelago and the castle also had a landed estate nearby with its own husbandry, farming and fishing (Kivikero 2020). There is also a great deal of correspondence from the noblemen to what is now mainland Sweden as well as to the nobility and priesthood of the cities Tallinn and Riga, capitals of Estonia and Latvia (Hausen 1934). Possibly sheep were imported to Åland from both sides of the Baltic Sea. The first sheep in the Åland Islands are dated to the Late Neolithic period, c. 3850-4400 cal BP (Storå 2000). Two right radii from Kastelholm were included in the study (Supplementary Table 6). The samples from Kastelholm castle were recovered in archaeological excavations in 1986 in an area outside and to the east of the Castle (Nordman 1991). The excavations revealed archaeological remains that were linked to four different chronological phases. The sampled sheep bones originate from the oldest layers, i.e. phase 1, which consist of a secondarily deposited landfill along the shore east of the castle. The archaeological finds show the date to fall around 1500-1550 AD, which is in good agreement with the radiocarbon dating of AKAS001 (Table 1).

### Radiocarbon dating

Samples were radiocarbon dated by the Tandem Laboratory at Uppsala University. Bone samples were mechanically cleaned by scraping and then ground in a mortar. 0.25 M HCl was added and incubated at ambient temperature for 48 hours. 0.01 M HCl was added to the insoluble fraction and incubated at 50 °C for 16 hours. The soluble fraction was added to a 30kDa ultrafilter and centrifuged. The retentate was lyophilized. Prior to accelerator determination the Fraction to be dated was combusted to CO2 using a Fe-catalyst. Acquired dates were calibrated using atmospheric data from IntCal20 (Reimer et al. 2020) in IOSACAL v.0.4.1 (Costa and Gutiérrez-Roig 2018).

### Sample preparation and sequencing

DNA was extracted from samples in a dedicated clean lab by the SciLifeLab aDNA unit at Uppsala University. Before sampling each bone was UV-irradiated (6 J/cm2 at 254 nm) on each side. The outer surface was then removed using a Dremel drill. Bones were wiped with first 0.5 % bleach and then Milli-Q water using sterile cotton swabs. A small piece of 50-100 mg was cut from each bone using a Dremel drill.

DNA was extracted using a modification to the protocol by (Yang et al. 1998), where 1 M urea replaced SDS. Extraction blanks were included as negative control. Samples were pre-treated with 1 ml of 0.5 M EDTA for 30 min at 37 °C. EDTA was removed, and samples digested with 1 ml of extraction buffer, under rotation at 37 °C for 22.5-23 h, and finally at 55 °C for 3.5-4 h. The supernatant was collected and concentrated using an Amicon Ultra-4 filter unit (Millipore). The extract was purified using a MinElute PCR purification kit (QIAGEN) according to standard instructions and finally eluted in 110 μl of EB buffer.

Libraries were prepared by the SciLifeLab aDNA unit at Uppsala University. Double stranded blunt end libraries were prepared according to (Meyer and Kircher 2010) with a MinElute PCR purification kit instead of SPRI beads. Libraries were quantified using qPCR in 25 μl reactions with Maxima SYBR green master mix (Thermo Fisher Scientific), 200 nM primer IS7 and 200 nM primer IS8 to determine the number of indexing cycles. Indexing was done in duplicates of 50 μl reactions using 6 μl of DNA library, 5 units of AmpliTaq Gold DNA polymerase (Thermo Fisher Scientific), 1’ GeneAmp Gold buffer (Thermo Fisher Scientific), 2.5 nM MgCL2, 250 μM of each dNTP, 200 nM of primer IS4 and 200 nM of indexing primer. PCR was performed as follows: 94 °C for 10 min, 14-17 cycles of (94 °C for 30 s, 60 °C for 30s, 72 °C for 45 s), and 72 °C for 10 min. Duplicates were pooled and purified with AMPure XP beads (Beckman Coulter). Quality of libraries were assessed using an Agilent 2200 Tapestation and Qubit dsDNA HS assay kits (Invitrogen).

Sequencing was done at the SciLifeLab SNP&SEQ Technology Platform at Uppsala University. Libraries were sequenced on an Illumina NovaSeq S4 flow cell using v1 chemistry and 100bp paired-end reads.

### Data preprocessing and mapping

Adapters were removed using CutAdapt v.2.3 (Martin 2011) using the following settings: --quality-base 33 --nextseq-trim=15 --overlap 3 -e 0.2 --trim-n --minimum-length 15:15. Reads were merged using FLASH v1.2.11 (Magoč and Salzberg 2011) with the following settings: --min-overlap 11 --max-overlap 150 --allow-outies. The quality of the reads was assessed using FastQC v0.11.9 (Andrews 2010). To corroborate that the sequences belonged to sheep, FastQScreen v.0.14.0 (Wingett and Andrews 2018) was run with a set of reference libraries composed of common contaminants plus those of a goat and a sheep.

Merged reads were mapped to the Texel sheep reference 3.1 (Oar3.1) using Bowtie2 v.2.3.5 (Langmead and Salzberg 2012) using the local alignment option, a mismatch penalty (--mp) of 4 and allowing for 1 mismatch in the seed alignment (-N 1). Other parameters were set as default. BAM files were sorted using SAMtools v.1.14 (Danecek et al. 2021) view and sort, respectively, using the standard settings. PCR duplicates were removed using DeDup v.0.12.8 (Peltzer et al. 2016). The merged reads were also mapped to the Texel sheep reference genome 4.0 (Oar4.0) (Consortium et al. 2010) containing the mitochondrial sequence from NCBI accession number NC_001941.1 using BWA aln v.0.7.17-r1188 (Li and Durbin 2009) and the non-default settings: -l 1024 -n 0.01 -o 2. Mapped reads were filtered using SAMtools v.1.14 (Danecek et al. 2021) view with the following settings: -q 30. These reads were sorted using SAMtools sort with standard settings. Duplicate reads were removed using DeDup v.0.12.8 (Peltzer et al. 2016) with standard settings. BAM files with duplicates removed were merged per individual using SAMtools merge and indexed using SAMtools index with standard settings for both Oar3.1 and Oar4.0. In order to validate the authenticity of the aDNA data, misincorporation patterns were calculated and visualized using mapDamage v2.0.9 (Jónsson et al. 2013), with standard settings. Overall mapping quality check was performed on both Oar3.1 and Oar4.0 with QualiMap v2.2.1 (Okonechnikov et al. 2016), and all quality control analyses were visualized using MultiQC v1.12 (Ewels et al. 2016).

### Molecular sex determination and mitochondrial haplotyping

Coverage was calculated using QualiMap v2.2.1 (Okonechnikov et al. 2016) on reads mapped to the modified Oar4.0 reference. Sex was determined by comparing the average coverage in the autosomes to the average coverage in the X chromosome using SAMtools depth, where a similar coverage was considered female and around half the coverage in the X chromosome was considered male.

Mapping Iterative Assembler (MIA) v.5a7fb5a (Stenzel 2013) was used to call consensus sequences for mitochondria in ancient samples. To remove reference bias, the Mouflon mitochondrial reference was used as well as a substitution matrix specific for aDNA. Consensus fasta-sequences were then assembled using sites with coverages higher than 3x, 15x and 50x as cutoff values, using Map Assember in MIA. As AKAS-001 has low coverage, a minimum 2x consensus was instead assembled for that sample.

As a measure of quality control, all mitogenome fasta files were aligned using MAFFT v.7.407 (Katoh and Standley 2013) with the standard settings, and inspected visually. For AKAS-001 the minimum 2x consensus fasta was chosen to call haplotypes, for AKAS-002 15x and for all ASTF samples the 50x. Haplotypes were assigned using MitoToolPy-Seq from MitoTool v.943ce25 (Peng et al. 2015) with settings: -s sheep - r whole.

### Reference data and genotyping

To obtain an unbiased genomic reference panel, all known Single Nucleotide Polymorphisms (SNP) from nine individuals of six different wild sheep species were collected from dbSNP (Supplementary Table 1). These SNPs were filtered to remove all multiallelic and duplicated SNPs and using the following settings in PLINK: --mind 0.1 --maf 0.05 --geno 0.1 --hwe 0.01. 188 published modern sheep genomes were selected from publications (Naval-Sanchez et al. 2018; Deng et al. 2020; Li et al. 2020) to cover a wide range of modern breeds (Supplementary Table 2). Raw sequencing reads for these individuals were downloaded and mapped with bwa mem (Li 2013). PCR duplicates were marked with Picard (http://broadinstitute.github.io/picard). SNPs were called and merged in these 188 sheep using GATK HaplotypeCaller (Van der Auwera and O’Connor 2020) using settings: --genotyping-mode GENOTYPE_GIVEN_ALLELES --output-mode EMIT_ALL_SITES. This set of SNPS was filtered using GATK VariantFiltration (Van der Auwera and O’Connor 2020) with the following filters: QD < 2.0, FS > 60.0, MQ < 40.0, SOR > 3.0, QUAL < 30.0, MQRankSum < -12.5, ReadPosRankSum < -8.0. This resulted in a reference panel of 2785430 SNPs. This reference panel was subset to include only Eurasian sheep breeds and Iranian Mouflons, and then filtered to remove low-quality SNPs, using PLINK v.1.90b4.9 (Purcell et al. 2007; Chang et al. 2015) with settings: --geno 0.01 --maf 0.01 --chr-set 26 no-y no-xy --allow-no-sex. Which left 161 modern individuals and 2100214 SNPs for analysis. We call this dataset WGS throughout the study.

To call SNPs in the ancient samples from sequences aligned to Oar 4.0, we used SAMtools mpileup v.1.14 (Danecek et al. 2021) with base alignment quality turned off and non-default settings: -B -q 30 -Q 30. At each position, a random read was drawn to code the individual as homozygous for that allele, at that position. A/G and C/T SNPs were coded as missing to minimize the effect of cytosine deamination.

To further situate the ancient samples in the Baltic Sea area a dataset consisting of European breeds genotyped on the Illumina Ovine Infinium® HD 600K chip was prepared(Supplementary Table 3). Data from (Rochus et al. 2018) was downloaded from Zenodo (Moreno-Romieux et al. 2017). Data from (Rochus et al. 2020a) was downloaded from DRYAD (Rochus et al. 2020b). Åland sheep data from (Cao et al. 2021) was acquired from the authors. Data for 16 Russian sheep breeds from (Igoshin et al. 2022) was downloaded from Figshare (Larkin and Zinovieva 2021). The reference datasets were merged and filtered with PLINK. First, duplicated positions and individuals were removed, and all heterozygous haploid variants were set to missing. Second, the dataset was filtered to remove low frequency SNPs with settings: --geno 0.01 --maf 0.01 --allow-extra-chr --allow-no-sex. This left 1010 individuals and 484 428 SNPs for analysis. SNPs for the ancient samples were again called as described above with the difference that reads aligned to Oar 3.1 were used. We call this dataset SNPCHP throughout the study.

As an outgroup, we also added pseudohaploid genotype calls from NGS data for 19 Iranian Asiatic Mouflon from the NextGen project (Ensembl 2022) which had been downloaded from the European Nucleotide Archive (ENA 2022). The genotype calls were performed the same way as for the ancient samples.

### Exploratory population genetic analysis

To prepare the dataset for PCA and ADMIXTURE data was pruned for SNPs in linkage disequilibrium (LD) to remove SNPs that were highly correlated, using PLINK with settings: --pairwise-indep 200 25 0.4 --chr-set 26 no-y no-xy --allow-no-sex. This left 578 285 SNPs of the WGS dataset and 275,544 SNPs of the SNPCHP dataset for analysis.

We run a Principal Component Analysis (PCA) to investigate clustering in the data, where ancient samples were projected on a reference panel, using smartpca v.16000 in eigensoft v.7.2.1 (Patterson et al. 2006; Price et al. 2006). To account for ancient individuals being possible outliers, as well as to deal with their amount of missing data these settings were used: altnormstyle: NO, numoutlieriter: 0, killr2: NO, numoutlierevec: 0, lsqproject: YES, shrinkmode: YES.

To further explore clustering with ADMIXTURE (Alexander and Lange 2011), the LD-pruned data set was converted to tped format using PLINK with settings: --recode transpose --chr-set 26 no-y no-xy --allow-no-sex and pseudo-haplodized by randomly choosing one allele at each site per individual in a tped file and coding the individual homozygous for that allele in that position. Pseudo-haplodizing the reference panel individuals reduces sample-specific drift in the ancient individuals that would solely be due to technical issues with the aDNA data. The pseudo-haplodized dataset was converted back to bed format using PLINK with settings: --make-bed --chr-set 26 no-y no-xy --allow-no-sex. ADMIXTURE v.1.3.0 (Alexander et al. 2009) was run, using standard settings and for K levels 2-10 with 20 different random seeds for each K.

### f statistics and diversity

Outgroup *f*_3_ statistics were calculated in ADMIXTOOLS v.2.0.0 (Maier et al. 2023) using the function qp3pop with standard settings and the non-pruned dataset described above without pre-calculating F2 statistics. The results were visualized using ggplot2 v.3.3.5 (Villanueva and Chen 2019) in R v.4.0.3 (R Core Team 2020). Conditional nucleotide diversity per group was calculated using POPSTATS (Skoglund et al. 2015), restricting the analysis to transversions (--notransitions) and allowing for more than 23 chromosome pairs (--not23).

Considering that the amount of sequencing data for ASTF-002 is relatively high for an ancient sample (11.6X), we decided to call diploid genotypes for this individual. We first realigned reads around indels using GATK (McKenna et al. 2010). Diploid genotypes were then called with the dedicated aDNA caller snpAD v0.3.4 (Prüfer 2018) restricting to reads with mapping qualities and base qualities of at least 30. Calls were performed separately for the BAM files aligned to Oar 3.1 and 4.0. Only sites with between 5 and 28 reads and genotype quality of at least 30 were considered for the analysis of runs-of-homozygosity (ROH). The diploid calls were merged with the modern reference data sets using plink excluding triallelic sites. ROH were then estimated using plink and the parameters --homozyg --homozyg-density 50, --homozyg-gap 1000, --homozyg-kb 500, --homozyg-snp 100, --homozyg-window-threshold 0.02, -- homozyg-window-snp 100, --homozyg-window-het 1, -- homozyg-window-missing 10, --allow-extra-chr, and --chr-set 26.

### Admixture graphs and continuity

We constructed admixture graphs from a reduced dataset to further investigate how ancient samples and NEST breeds are related. The SNPCHP data set was subset to include only Mouflons, Swedish NEST breeds, and Russian breeds. 502 individuals remained for the analysis. The data set was converted to VCF using PLINK and all sites with missing calls were removed with settings: --allow-no-sex --allow-extra-chr -- recode vcf-iid –geno 0 and all modern genomes were pseudo-haploidized to match the wild mouflons and the ancient samples. Then the VCF was in turn converted to .tmix format using vcf2treemix.py (Silva 2017). The dataset was then run on OrientAGraph v1.0 (Molloy et al. 2021), with a fixed seed to retain the same starting tree structure for each run, for migration edges 0-3 (Supplementary Figure 7), chosen as three edges allows for reasonable running time, using settings: -seed 32295 -mlno-allmigs and setting the the Asiatic Mouflons as the root of the tree. Trees and residuals were visualized in R v.4.0.3 (R Core Team 2020) using the packages RColorBrewer (Neuwirth 2014) and R.utils (Bengtsson 2021) with the functions plot_tree provided in TreeMix (Pickrell and Pritchard 2012) (Supplementary Figure 7).

As a complementary approach, we employed qpGraph (Patterson et al. 2012) to reconstruct admixture graphs. We used the find_graphs function in ADMIXTOOLS v.2.0.0 (Maier et al. 2023) to explore the graph space. *f2* statistics were calculated using f2_from_geno allowing for 15% missingness per SNP and more than 22 chromosome pairs. find_graphs was then run with Asiatic mouflon as an outgroup for 1000 different random seeds and the parameters stop_gen = 10000, stop_gen2 = 30 and plusminus_generations = 10. Later, we also constrained the search space by forbidding admixture into the ancient groups as well as the European mouflons as a representative of the Mediterranean Neolithic expansion. For these runs, we first generated random graphs using random_admixturegraph and ntry = 10000 in order to start with a graph that fulfills the constraints. The randomly generated graph was then used as initgraph for find_graphs. We also explored 1000 different random seeds for the constrained graphs. For both options we explored between 0 and 5 migration events. Graph fits were compared using the qpgraph_resample_multi and compare_fits functions.

In order to test if there has been direct population continuity within the Baltic sheep population through the time period covered by our samples, we used a maximum likelihood ratio test described in (Schraiber 2018) (https://github.com/Schraiber/continuity). We used this explicit test for population continuity both on NEST and non-NEST breeds with 3 or more samples from the WGS and the SNP-CHP datasets. In order to polarize the ancestral and derived alleles of our modern datasets, we used a Caphi goat (ENA accession ERR219543) and an Inner Mongolia cashmere goat (SRA accession SRR5557418) mapped to Oar4.0 and Oar3.1, respectively, as the outgroup species. Homozygous alleles in the goat were assigned as ancestral variants. Data was then filtered for high and low coverage sites with the default cutoffs and then two models were fitted to each dataset: one assuming continuity and the other assuming independent populations. The resulting likelihoods were then used to calculate the likelihood ratio statistic.

### Phenotypic analysis

To investigate if the ancient samples in this study had or lacked the insertion polymorphism causative of a fine wool phenotype, reads were mapped to two modified Oar3.1 references. One containing the full insertion and one lacking the full insertion according to (Demars et al. 2017; Rossi et al. 2021). Merged reads from the samples were mapped with BWA aln v.0.7.17-r1188 (Li and Durbin 2009). These reads were sorted using SAMtools v.1.14 (Danecek et al. 2021) sort with standard settings. Duplicate reads were removed using SAMtools markdup. Samples were then inspected visually in IGV v.2.11.4 (Robinson et al. 2011) to find out how many reads crossed the breakpoints of the insertion and the point in the genome lacking the insertion where the insertion would be situated, according to (Rossi et al. 2021).

Additionally, the alleles of ancient samples at SNP rs161553028, which is associated with fiber diameter (Lv et al. 2022), were visually inspected in IGV v.2.11.4 (Robinson et al. 2011) in the reads aligned to Oar3.1.

## Data availability

Raw sequence data and aligned reads for the five new ancient individuals are available through the European Nucleotide Archive under accession number PRJEB59481. Genotype calls for the ancient individuals are made available on Zenodo (10.5281/zenodo.7862142).

## Acknowledgements

We thank Conor Rossi and Kevin Daly for discussion on genotyping the insertion and for sharing modified reference genomes. We thank the sheep research community and specifically the authors of our sources for making their genotype and sequence data available to the public. This work was supported by a grant from the Swedish Research Council Vetenskapsrådet (2017–05267) to TG. TG was partly funded through a Riksbankens Jubileumsfond grant (P21-0266) awarded to Helena Malmström. AERS was supported by a postdoctoral stipend from Carl Tryggers Stiftelse för Vetenskaplig Forskning (CTS 18:129). Processing of aDNA was performed by the SciLifeLab Ancient DNA facility. Sequencing was performed by the SNP&SEQ Technology Platform in Uppsala, part of the National Genomics Infrastructure (NGI) Sweden and Science for Life Laboratory. The SNP&SEQ Platform is also supported by the Swedish Research Council and the Knut and Alice Wallenberg Foundation. The computations and data handling were enabled by resources in projects SNIC 2021/22-705, SNIC 2021/22-763, SNIC 2021/23-637, SNIC 2021/2-17 and SNIC 2022/2-11 provided by the National Academic Infrastructure for Supercomputing in Sweden (NAISS) and the Swedish National Infrastructure for Computing (SNIC) at Uppmax, partially funded by the Swedish Research Council through grant agreements no. 2022-06725 and no. 2018-05973.

## References

Alexander DH, Lange K. 2011. Enhancements to the ADMIXTURE algorithm for individual ancestry estimation. BMC Bioinformatics 12:246.

Alexander DH, Novembre J, Lange K. 2009. Fast model-based estimation of ancestry in unrelated individuals. Genome Res. 19:1655–1664.

Allentoft ME, Sikora M, Sjögren K-G, Rasmussen S, Rasmussen M, Stenderup J, Damgaard PB, Schroeder H, Ahlström T, Vinner L, et al. 2015. Population genomics of Bronze Age Eurasia. Nature 522:167–172.

Andrews S. 2010. FastQC A Quality Control tool for High Throughput Sequence Data. Available from: https://www.bioinformatics.babraham.ac.uk/projects/fastqc/

Apel J, Wallin P, Storå J, Possnert G. 2018. Early Holocene human population events on the island of Gotland in the Baltic Sea (9200-3800 cal. BP). Quaternary International 465:276–286.

Bengtsson H. 2021. R.utils: Various Programming Utilities. Available from: https://CRAN.R-project.org/package=R.utils

Cao Y-H, Xu S-S, Shen M, Chen Z-H, Gao L, Lv F-H, Xie X-L, Wang X-H, Yang H, Liu C-B, et al. 2021. Historical Introgression from Wild Relatives Enhanced Climatic Adaptation and Resistance to Pneumonia in Sheep. Molecular Biology and Evolution 38:838–855.

Chang CC, Chow CC, Tellier LC, Vattikuti S, Purcell SM, Lee JJ. 2015. Second-generation PLINK: rising to the challenge of larger and richer datasets. Gigascience 4:s13742–015.

Chessa B, Pereira F, Arnaud F, Amorim A, Goyache F, Mainland I, Kao RR, Pemberton JM, Beraldi D, Stear MJ, et al. 2009. Revealing the history of sheep domestication using retrovirus integrations. Science 324:532–536.

Consortium TISG, Archibald AL, Cockett NE, Dalrymple BP, Faraut T, Kijas JW, Maddox JF, McEwan JC, Hutton Oddy V, Raadsma HW, et al. 2010. The sheep genome reference sequence: a work in progress. Animal Genetics 41:449–453.

Costa S, Gutiérrez-Roig M. 2018. IOSACal. Available from: https://doi.org/10.5281/zenodo.1243291

Coutinho A, Günther T, Munters AR, Svensson EM, Götherström A, Storå J, Malmström H, Jakobsson M. 2020. The Neolithic Pitted Ware culture foragers were culturally but not genetically influenced by the Battle Axe culture herders. American Journal of Physical Anthropology 172:638–649.

Danecek P, Bonfield JK, Liddle J, Marshall J, Ohan V, Pollard MO, Whitwham A, Keane T, McCarthy SA, Davies RM. 2021. Twelve years of SAMtools and BCFtools. Gigascience 10:giab008.

Demars J, Cano M, Drouilhet L, Plisson-Petit F, Bardou P, Fabre S, Servin B, Sarry J, Woloszyn F, Mulsant P, et al. 2017. Genome-wide identification of the mutation underlying fleece variation and discriminating ancestral hairy species from modern woolly sheep. Molecular biology and evolution 34:1722–1729.

Deng J, Xie X-L, Wang D-F, Zhao C, Lv F-H, Li X, Yang J, Yu J-L, Shen M, Gao L. 2020. Paternal origins and migratory episodes of domestic sheep. Current Biology 30:4085–4095.

Dýrmundsson ÓR, Niżnikowski R. 2010. North European short-tailed breeds of sheep: a review. Animal 4:1275–1282.

Edberg R. 1986. Gutefåren och deras ursprung. Fataburen:151–162.

ENA. 2022. Project: PRJEB3139. European Nucleotide Archive [Internet]. Available from: https://www.ebi.ac.uk/ena/browser/view/PRJEB3139?show=reads

Ensembl. 2022. NextGen. NextGen [Internet]. Available from: https://projects.ensembl.org/nextgen/

Ewels P, Magnusson M, Lundin S, Käller M. 2016. MultiQC: summarize analysis results for multiple tools and samples in a single report. Bioinformatics 32:3047–3048.

Fages A, Hanghøj K, Khan N, Gaunitz C, Seguin-Orlando A, Leonardi M, McCrory Constantz C, Gamba C, Al-Rasheid KAS, Albizuri S, et al. 2019. Tracking Five Millennia of Horse Management with Extensive Ancient Genome Time Series. Cell 177:1419–1435.e31.

Fischer A. 2002. Food for feasting. The neolithisation of Denmark 150:343–393.

Frantz LAF, Bradley DG, Larson G, Orlando L. 2020. Animal domestication in the era of ancient genomics. Nat Rev Genet:1–12.

Ghoreishifar SM, Rochus CM, Moghaddaszadeh-Ahrabi S, Davoudi P, Salek Ardestani S, Zinovieva NA, Deniskova TE, Johansson AM. 2021. Shared Ancestry and Signatures of Recent Selection in Gotland Sheep. Genes 12:433.

Günther T, Malmström H, Svensson EM, Omrak A, Sánchez-Quinto F, Kılınç GM, Krzewińska M, Eriksson G, Fraser M, Edlund H, et al. 2018. Population genomics of Mesolithic Scandinavia: Investigating early postglacial migration routes and high-latitude adaptation. PLOS Biology 16:e2003703.

Haak W, Lazaridis I, Patterson N, Rohland N, Mallick S, Llamas B, Brandt G, Nordenfelt S, Harney E, Stewardson K, et al. 2015. Massive migration from the steppe was a source for Indo-European languages in Europe. Nature [Internet]. Available from: http://www.nature.com/doifinder/10.1038/nature14317

Hausen R. 1934. Kastelholms slott och dess borgherrar. Helsinki: Swedish Literary Society in Finland

Igoshin A, Deniskova T, Yurchenko A, Yudin N, Dotsev A, Selionova M, Zinovieva N, Larkin D. 2022. Copy number variants in genomes of local sheep breeds from Russia. Animal genetics.

Jónsson H, Ginolhac A, Schubert M, Johnson PL, Orlando L. 2013. mapDamage2. 0: fast approximate Bayesian estimates of ancient DNA damage parameters. Bioinformatics 29:1682–1684.

Katoh K, Standley DM. 2013. MAFFT multiple sequence alignment software version 7: improvements in performance and usability. Molecular biology and evolution 30:772–780.

Kivikero H. 2020. The Economy of Food : Tracing food production and consumption in the Castles of Kastelholm and Raseborg from the 14th to 16th centuries. Available from: https://helda.helsinki.fi/handle/10138/313968

Langmead B, Salzberg SL. 2012. Fast gapped-read alignment with Bowtie 2. Nat Methods 9:357–359.

Larkin D, Zinovieva N. 2021. Genotyping data for 16 Russian sheep breeds. Available from: https://figshare.com/articles/dataset/Genotyping_data_for_16_Russian_sheep_breeds/16806781

Li H. 2013. Aligning sequence reads, clone sequences and assembly contigs with BWA-MEM. Available from: http://arxiv.org/abs/1303.3997

Li H, Durbin R. 2009. Fast and accurate short read alignment with Burrows–Wheeler transform. bioinformatics 25:1754–1760.

Li X, Yang J, Shen M, Xie X-L, Liu G-J, Xu Y-X, Lv F-H, Yang H, Yang Y-L, Liu C-B, et al. 2020. Whole-genome resequencing of wild and domestic sheep identifies genes associated with morphological and agronomic traits. Nature Communications 11:2815.

Librado P, Khan N, Fages A, Kusliy MA, Suchan T, Tonasso-Calvière L, Schiavinato S, Alioglu D, Fromentier A, Perdereau A, et al. 2021. The origins and spread of domestic horses from the Western Eurasian steppes. Nature 598:634–640.

Lindqvist C, Possnert G. 1997. The subsistence economy and diet at Jakobs/Ajvide, Eksta parish and other prehistoric dwelling and burial sites on Gotland in longterm perspective. In: Burenhult G, editor. Remote sensing. Vol. 1. p. 29–90.

Lindqvist C, Possnert G. 1999. The first Seal Hunter Families on Gotland. Current Swedish Archaeology 7:65.

Lv F-H, Cao Y-H, Liu G-J, Luo L-Y, Lu R, Liu M-J, Li W-R, Zhou P, Wang X-H, Shen M, et al. 2022. Whole-Genome Resequencing of Worldwide Wild and Domestic Sheep Elucidates Genetic Diversity, Introgression, and Agronomically Important Loci. Molecular Biology and Evolution 39:msab353.

Machová K, Málková A, Vostrý L. 2022. Sheep Post-Domestication Expansion in the Context of Mitochondrial and Y Chromosome Haplogroups and Haplotypes. Genes 13:613.

Magoč T, Salzberg SL. 2011. FLASH: fast length adjustment of short reads to improve genome assemblies. Bioinformatics 27:2957–2963.

Maier R, Flegontov P, Flegontova O, Isildak U, Changmai P, Reich D. 2023. On the limits of fitting complex models of population history to f-statistics.Nordborg M, editor. eLife 12:e85492.

Malmer MP. 2002. The Neolithic of south Sweden. 1st ed. Stockholm: Royal Swedish Academy of Letters, History and Antiquities

Malmström H, Günther T, Svensson EM, Juras A, Fraser M, Munters AR, Pospieszny Ł, Tõrv M, Lindström J, Götherström A. 2019. The genomic ancestry of the Scandinavian Battle Axe Culture people and their relation to the broader Corded Ware horizon. Proceedings of the Royal Society B 286:20191528.

Marciniak A. 2011. The Secondary Products Revolution: Empirical Evidence and its Current Zooarchaeological Critique. J World Prehist 24:117–130.

Martin M. 2011. Cutadapt removes adapter sequences from high-throughput sequencing reads. EMBnet.journal 17:10–12.

McKenna A, Hanna M, Banks E, Sivachenko A, Cibulskis K, Kernytsky A, Garimella K, Altshuler D, Gabriel S, Daly M, et al. 2010. The Genome Analysis Toolkit: A MapReduce framework for analyzing next-generation DNA sequencing data. Genome Res 20:1297–1303.

Meyer M, Kircher M. 2010. Illumina Sequencing Library Preparation for Highly Multiplexed Target Capture and Sequencing. Cold Spring Harbor Protocols 2010:pdb.prot5448-pdb.prot5448.

Molloy EK, Durvasula A, Sankararaman S. 2021. Advancing admixture graph estimation via maximum likelihood network orientation. Bioinformatics 37:i142–i150.

Moreno-Romieux C, Tortereau F, Raoul J, Bertrand S. 2017. High density genotypes of French Sheep populations. Zenodo [Internet]. Available from: https://doi.org/10.5281/zenodo.237116

Mourier T, Ho SY, Gilbert MTP, Willerslev E, Orlando L. 2012. Statistical guidelines for detecting past population shifts using ancient DNA. Molecular biology and evolution 29:2241–2251.

Naval-Sanchez M, Nguyen Q, McWilliam S, Porto-Neto LR, Tellam R, Vuocolo T, Reverter A, Perez-Enciso M, Brauning R, Clarke S, et al. 2018. Sheep genome functional annotation reveals proximal regulatory elements contributed to the evolution of modern breeds. Nature communications 9:859.

Neuwirth E. 2014. RColorBrewer: ColorBrewer Palettes. Available from: https://CRAN.R-project.org/package=RColorBrewer

Niemi M, Bläuer A, Iso-Touru T, Nyström V, Harjula J, Taavitsainen J-P, Stor\a a J, Lidén K, Kantanen J. 2013. Mitochondrial DNA and Y-chromosomal diversity in ancient populations of domestic sheep (Ovis aries) in Finland: comparison with contemporary sheep breeds. Genetics Selection Evolution 45:2.

Niemi M, Sajantila A, Ahola V, Vilkki J. 2018. Sheep and cattle population dynamics based on ancient and modern DNA reflects key events in the human history of the North-East Baltic Sea Region. Journal of Archaeological Science: Reports 18:169–173.

Nordman A-M. 1991. Arkeologisk undersökning 12.23 Kastelholm; Kastelholms slott KS 41 - VVS öster om slottet. In: Kastelholms slott Arkeologiska undersökningar 1985-1989; KS 30-KS 52. Mariehamn: Ålands landskapsstyrelse, Museibyrån. p. 339–362.

Okonechnikov K, Conesa A, García-Alcalde F. 2016. Qualimap 2: advanced multisample quality control for high-throughput sequencing data. Bioinformatics 32:292–294.

Palamarz P. 2004. Kastelholms slott : från medeltida borg till byggnadsminne. [Mariehamn]: Ålands landskapsstyrelse, Museibyrån

Patterson N, Moorjani P, Luo Y, Mallick S, Rohland N, Zhan Y, Genschoreck T, Webster T, Reich D. 2012. Ancient Admixture in Human History. Genetics 192:1065–1093.

Patterson N, Price AL, Reich D. 2006. Population Structure and Eigenanalysis. PLoS Genetics 2:e190.

Peltzer A, Jäger G, Herbig A, Seitz A, Kniep C, Krause J, Nieselt K. 2016. EAGER: efficient ancient genome reconstruction. Genome biology 17:1–14.

Peng M-S, Fan L, Shi N-N, Ning T, Yao Y-G, Murphy RW, Wang W-Z, Zhang Y-P. 2015. DomeTree: a canonical toolkit for mitochondrial DNA analyses in domesticated animals. Molecular Ecology Resources 15:1238–1242.

Pickrell JK, Pritchard JK. 2012. Inference of population splits and mixtures from genome-wide allele frequency data. PLoS Genet. 8:e1002967.

Price AL, Patterson NJ, Plenge RM, Weinblatt ME, Shadick NA, Reich D. 2006. Principal components analysis corrects for stratification in genome-wide association studies. Nat Genet 38:904–909.

Prüfer K. 2018. snpAD: an ancient DNA genotype caller. Bioinformatics 34:4165–4171.

Purcell S, Neale B, Todd-Brown K, Thomas L, Ferreira MAR, Bender D, Maller J, Sklar P, de Bakker PIW, Daly MJ, et al. 2007. PLINK: a tool set for wholegenome association and population-based linkage analyses. Am. J. Hum. Genet. 81:559–575.

R Core Team. 2020. R: A Language and Environment for Statistical Computing. Available from: https://www.R-project.org/

Rannamäe E, Lõugas L, Niemi M, Kantanen J, Maldre L, Kadõrova N, Saarma U. 2016a. Maternal and paternal genetic diversity of ancient sheep in Estonia from the Late Bronze Age to the post-medieval period and comparison with other regions in Eurasia. Animal genetics.

Rannamäe E, Lõugas L, Speller CF, Valk H, Maldre L, Wilczyński J, Mikhailov A, Saarma U. 2016b. Three thousand years of continuity in the maternal lineages of ancient sheep (Ovis aries) in Estonia. PloS one 11:e0163676.

Rannamäe E, Saarma U, Ärmpalu-Idvand A, Teasdale MD, Speller C. 2020. Retroviral analysis reveals the ancient origin of Kihnu native sheep in Estonia: implications for breed conservation. Scientific reports 10:1–8.

Reimer PJ, Austin WE, Bard E, Bayliss A, Blackwell PG, Ramsey CB, Butzin M, Cheng H, Edwards RL, Friedrich M. 2020. The IntCal20 Northern Hemisphere radiocarbon age calibration curve (0–55 cal kBP). Radiocarbon 62:725–757.

Robinson JT, Thorvaldsdóttir H, Winckler W, Guttman M, Lander ES, Getz G, Mesirov JP. 2011. Integrative genomics viewer. Nature biotechnology 29:24–26.

Rochus CM, Jonas E, Johansson AM. 2020a. Population structure of five native sheep breeds of Sweden estimated with high density SNP genotypes. BMC genetics 21:1–9.

Rochus CM, Jonas E, Johansson AM. 2020b. Population structure of five native sheep breeds of Sweden estimated with high density SNP genotypes. DRYAD [Internet]. Available from: https://doi.org/10.5061/dryad.34tmpg4gj

Rochus CM, Tortereau F, Plisson-Petit F, Restoux G, Moreno-Romieux C, Tosser-Klopp G, Servin B. 2018. Revealing the selection history of adaptive loci using genome-wide scans for selection: an example from domestic sheep. BMC genomics 19:1–17.

Rossi C, Ruß-Popa G, Mattiangeli V, McDaid F, Hare AJ, Davoudi H, Laleh H, Lorzadeh Z, Khazaeli R, Fathi H. 2021. Exceptional ancient DNA preservation and fibre remains of a Sasanian saltmine sheep mummy in Chehrābād, Iran. Biology Letters 17:20210222.

Rundkvist M, Lindqvist C, Thorsberg K. 2004. Barshalder 3 Rojrhage in Grötlingbo : a multi-component Neolithic shore site on Gotland. Stockholm: Dept. of Archaeology [Institutionen för arkeologi], Univ. Available from: http://bibliotek.gotland.se/index2.nsf/0/0fbac78c5887de8cc1256d4200411f96/$FILE/bhr2.pdf

Ryder ML. 1983. Sheep & man. London: Duckworth

Ryder ML. 1984. Sheep. In: Mason IL, editor. Evolution of domesticated animals. London, England: Longman. p. 63–84.

Schnittger B, Rydh H. 1940. Grottan Stora Förvar på Stora Karlsö. Stockholm: Kungl. Vitterhets Historie och Antikvitets akademien

Schraiber JG. 2018. Assessing the relationship of ancient and modern populations. Genetics 208:383–398.

Sherratt A. 1983. The secondary exploitation of animals in the Old World. World Archaeology 15:90–104.

Silva D. 2017. vcf2treemix.py. Available from: https://github.com/CoBiG2/RAD_Tools/blob/master/vcf2treemix.py

Sjögren KG, Axelson T, Vretemark M. 2019. Middle Neolithic economy in Falbygden, Sweden Preliminary results from Karleby Logården. In: International Conference Megaliths, Societies, Landscapes. Early Monumentality and Social Differentiation in Neolithic Europe (2015. Kiel) Megaliths, societies, landscapes: early monumentality and social differentiation in Neolithic Europe : proceedings of the international conference 16th-20th june 2015 in Kiel,Vol. 2, Tomo 2, 2019 (Volume 2), ISBN 978-3-7749-4213-4, págs. 705-718. Verlag Dr. Rudolf Habelt GmbH. p. 705–718. Available from: https://dialnet.unirioja.es/servlet/articulo?codigo=8420261

Skoglund P, Mallick S, Bortolini MC, Chennagiri N, Hünemeier T, Petzl-Erler ML, Salzano FM, Patterson N, Reich D. 2015. Genetic evidence for two founding populations of the Americas. Nature 525:104–108.

Skoglund P, Malmstrom H, Omrak A, Raghavan M, Valdiosera C, Gunther T, Hall P, Tambets K, Parik J, Sjogren K-G, et al. 2014. Genomic Diversity and Admixture Differs for Stone-Age Scandinavian Foragers and Farmers. Science 344:747–750.

Smith O, Nicholson WV, Kistler L, Mace E, Clapham A, Rose P, Stevens C, Ware R, Samavedam S, Barker G. 2019. A domestication history of dynamic adaptation and genomic deterioration in Sorghum. Nature plants 5:369–379.

Stenzel U. 2013. mapping-iterative-assembler. Available from: https://github.com/mpieva/mapping-iterative-assembler

Stoffel MA, Johnston SE, Pilkington JG, Pemberton JM. 2021. Genetic architecture and lifetime dynamics of inbreeding depression in a wild mammal. Nat Commun 12:2972.

Storå J. 2000. Sealing and animal husbandry in the Ålandic Middle and Late Neolithic. Fennoscandia Archaeologica [Internet]. Available from: https://journal.fi/fennoscandiaarchaeologica/article/view/126697

Swedish Board of Agriculture. 2022. Statistik över svenska husdjursraser. Jordbruksverket [Internet]. Available from: https://jordbruksverket.se/djur/lantbruksdjur-och-hastar/husdjursraser-och-avelsorganisationer/forskning-och-aktiviteter-om-husdjursraser/arkiv/2022-03-17-statistik-over-svenska-husdjursraser

Tapio M, Tapio I, Grislis Z, Holm L-E, Jeppsson S, Kantanen J, Miceikiene I, Olsaker I, Viinalass H, Eythorsdottir E. 2005. Native breeds demonstrate high contributions to the molecular variation in northern European sheep. Molecular Ecology 14:3951–3963.

Taylor WT, Pruvost M, Posth C, Rendu W, Krajcarz MT, Abdykanova A, Brancaleoni G, Spengler R, Hermes T, Schiavinato S. 2021. Evidence for early dispersal of domestic sheep into Central Asia. Nature Human Behaviour:1–11.

Van der Auwera GA, O’Connor BD. 2020. Genomics in the cloud: using Docker, GATK, and WDL in Terra. O’Reilly Media

Villanueva RAM, Chen ZJ. 2019. ggplot2: elegant graphics for data analysis.

Wingett SW, Andrews S. 2018. FastQ Screen: A tool for multi-genome mapping and quality control. Available from: https://f1000research.com/articles/7-1338

Yang DY, Eng B, Waye JS, Dudar JC, Saunders SR. 1998. Technical note: improved DNA extraction from ancient bones using silica-based spin columns. Am. J. Phys. Anthropol. 105:539–543.

Yurtman E, Özer O, Yüncü E, Dağtaş ND, Koptekin D, Çakan YG, Özkan M, Akbaba A, Kaptan D, Atağ G, et al. 2021. Archaeogenetic analysis of Neolithic sheep from Anatolia suggests a complex demographic history since domestication. Commun Biol 4:1–11.

Zeder MA. 2008. Domestication and early agriculture in the Mediterranean Basin: Origins, diffusion, and impact. Proceedings of the national Academy of Sciences 105:11597–11604.

