## Supplementary Figures for "Ancient Sheep Genomes reveal four Millennia of North European Short-Tailed Sheep in the Baltic Sea region"

Martin NA Larsson[1,†], Pedro Morell Miranda[1,†], Li Pan[1], Kıvılcım Başak Vural[2], Damla Kaptan[2], André Elias Rodrigues Soares[1], Hanna Kivikero[3], Juha Kantanen[4], Mehmet Somel[2], Füsun Özer[5], Anna M Johansson[6], Jan Storå[7], Torsten Günther[1,*]

1 Human Evolution, Department of Organismal Biology, Uppsala University, Sweden
2 Department of Biological Sciences, Middle East Technical University, Ankara, Turkey
3 Department of Culture, University of Helsinki, Helsinki, Finland
4 Natural Resources Institute Finland, Jokioinen, Finland
5 Department of Anthropology, Hacettepe University, Ankara, Turkey
6 Department of Animal Breeding and Genetics, Swedish University of Agricultural Sciences, Uppsala, Sweden
7 Osteological Research Laboratory, University of Stockholm, Stockholm, Sweden

**AKAS-001** out.extendedFrags.fastq.sorted

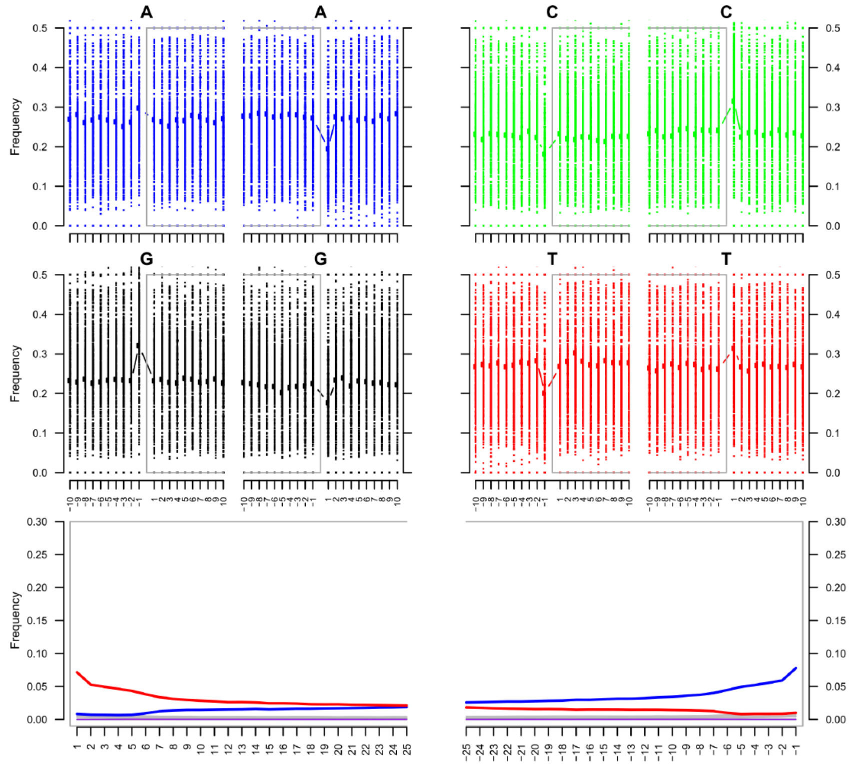

**AKAS-002** out.extendedFrags.fastq.sorted

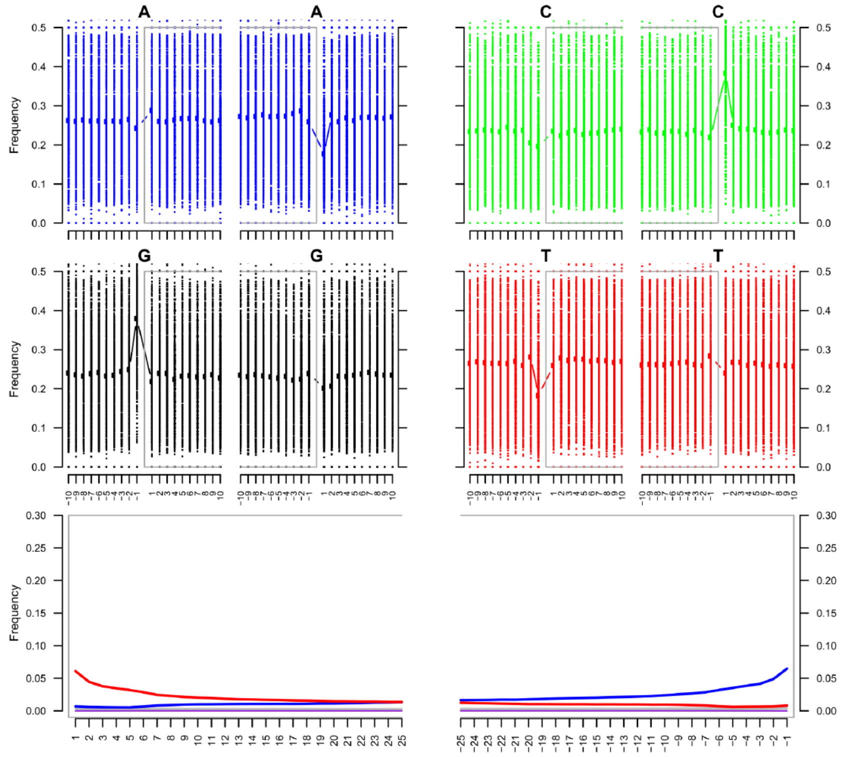


**ASTF-001** out.extendedFrags.fastq.sorted

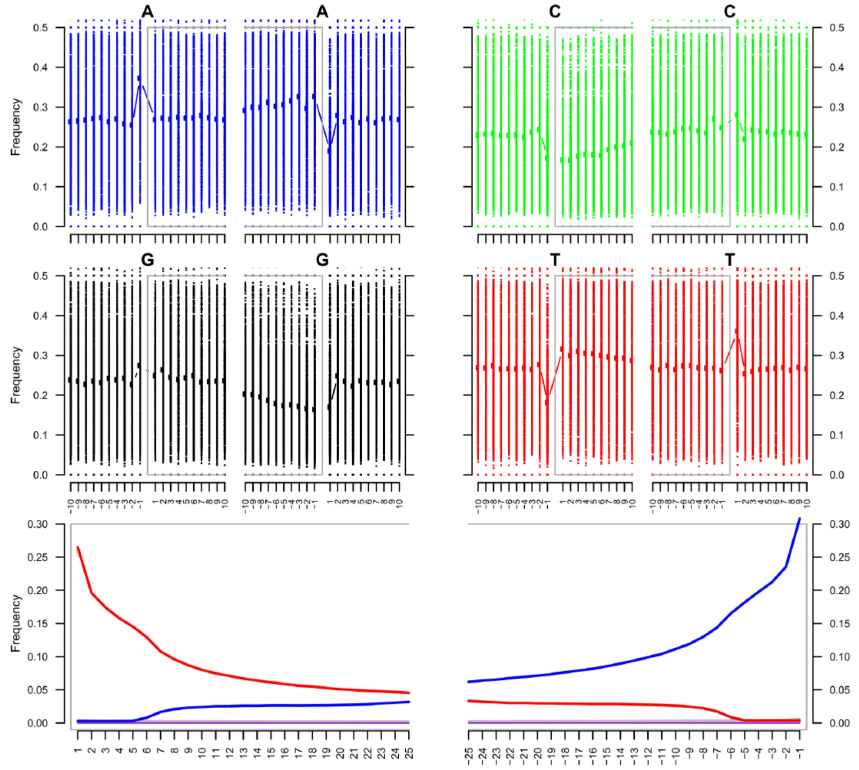

**ASTF-002** out.extendedFrags.fastq.sorted

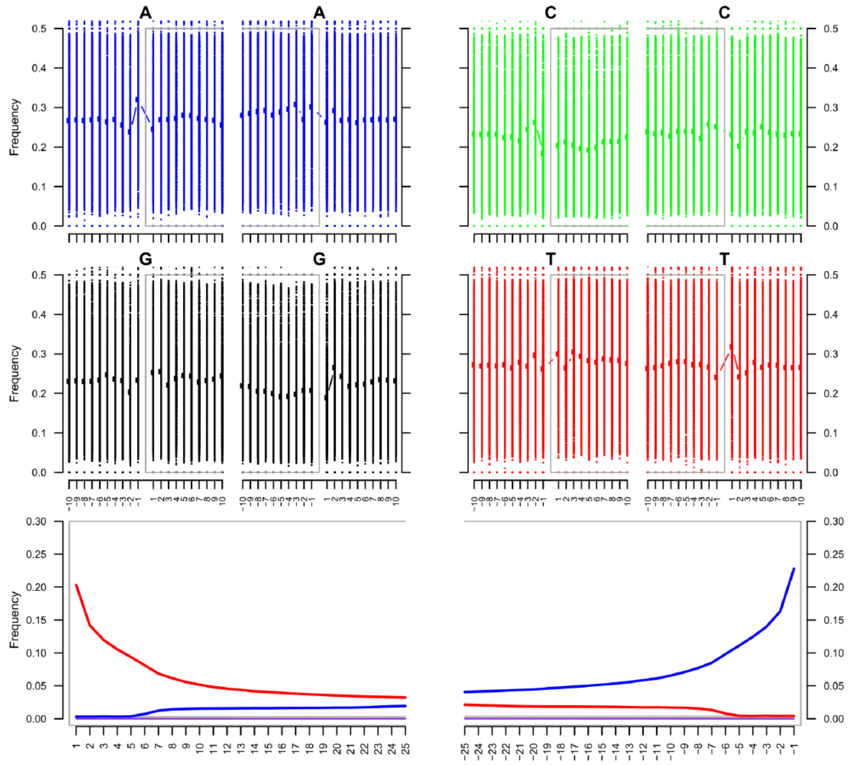


**ASTF-003** out.extendedFrags.fastq.sorted

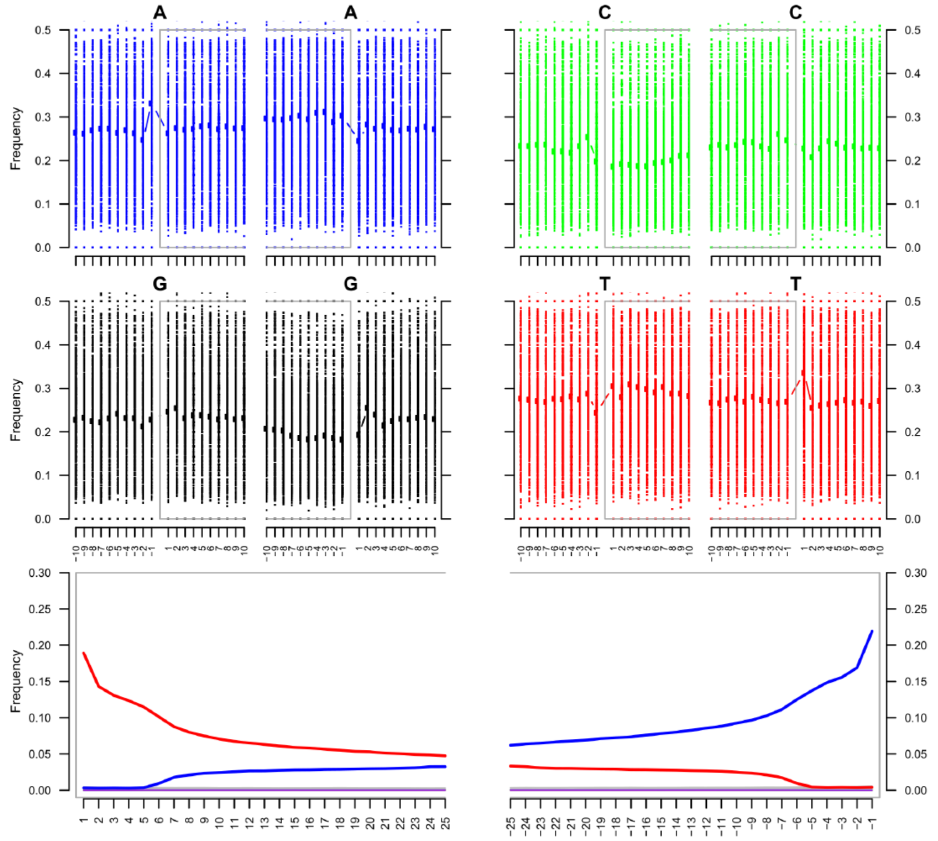


**Supplementary Figure 1**: mapDamage misincorporation plots for the 5 ancient samples.

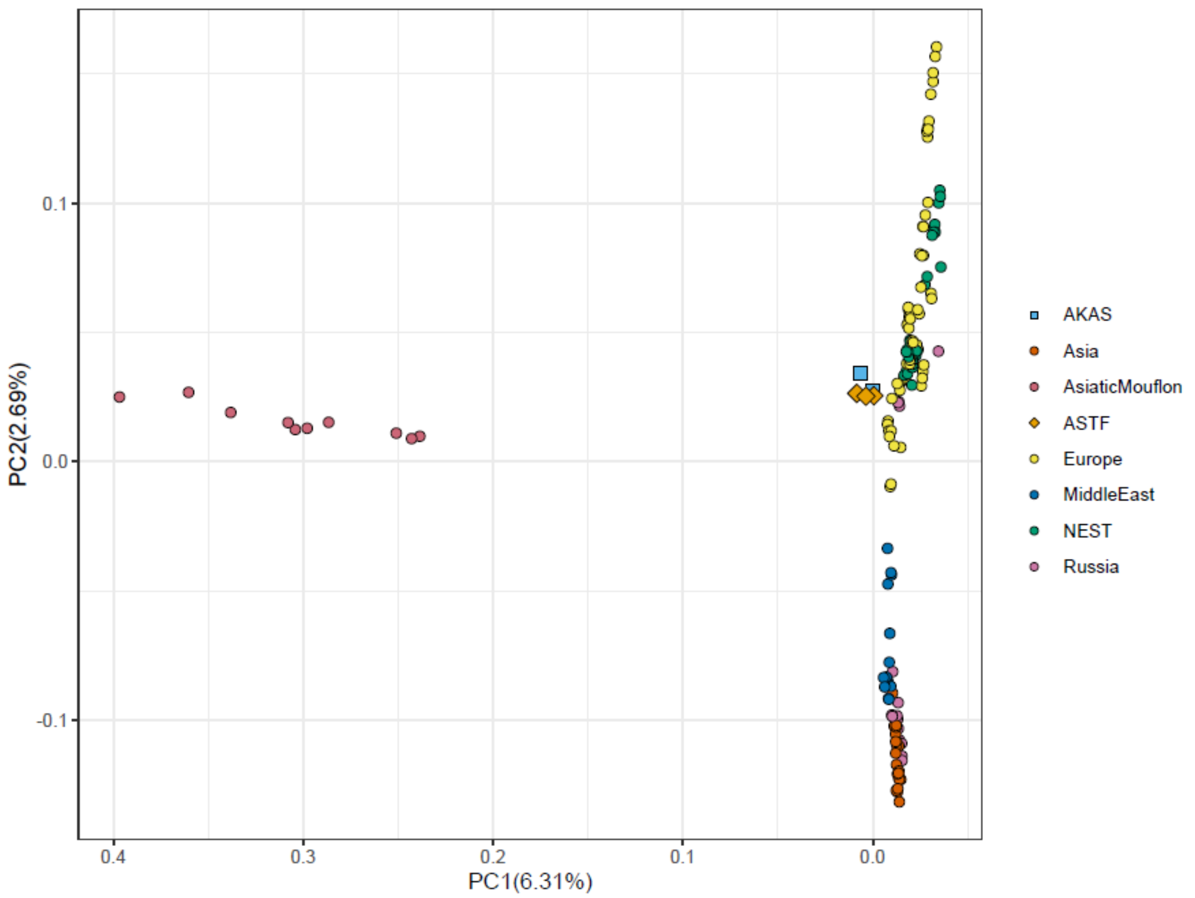


**Supplementary Figure 2**: WGS PCA showing PC 1 vs PC 2, percentage of explained variation is shown within parentheses of axis titles.

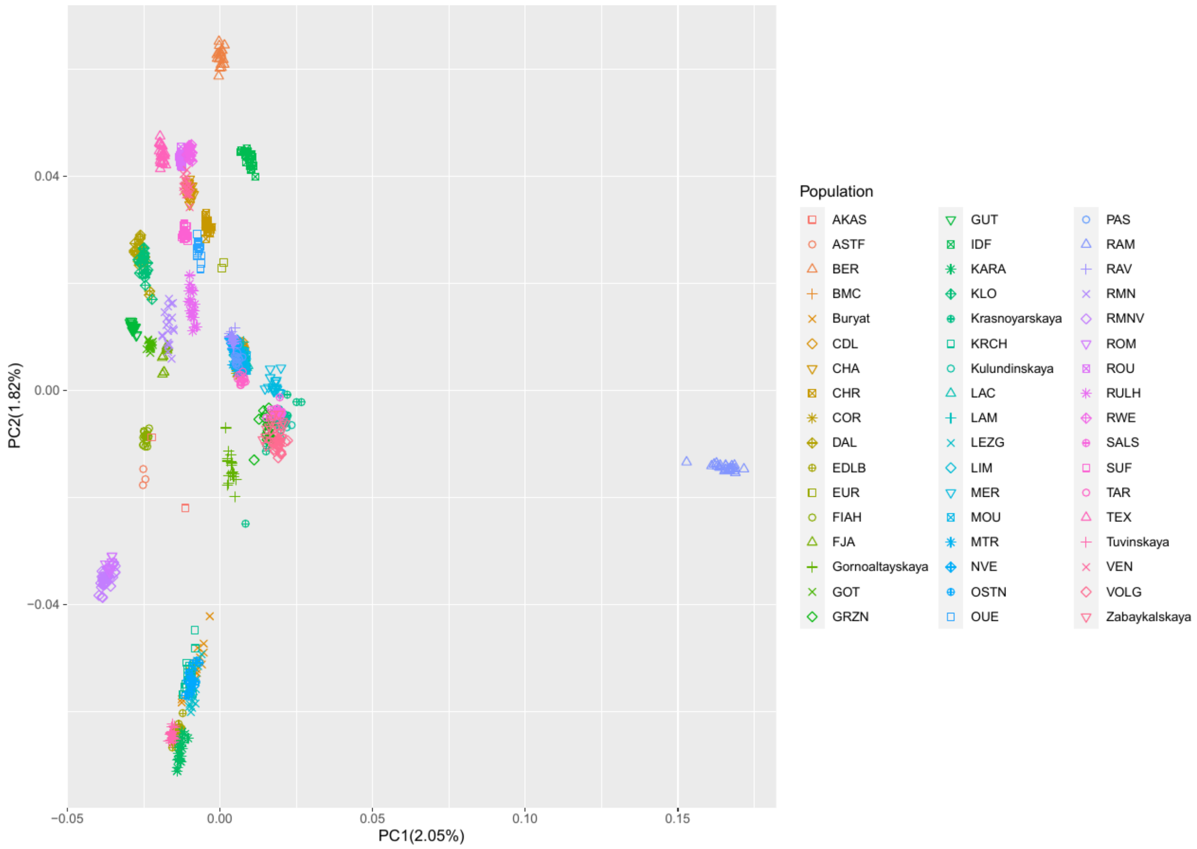


**Supplementary Figure 3**: SNPCHP PCA showing PC 1 vs PC 2, percentage of explained variation is shown within parentheses of axis titles.

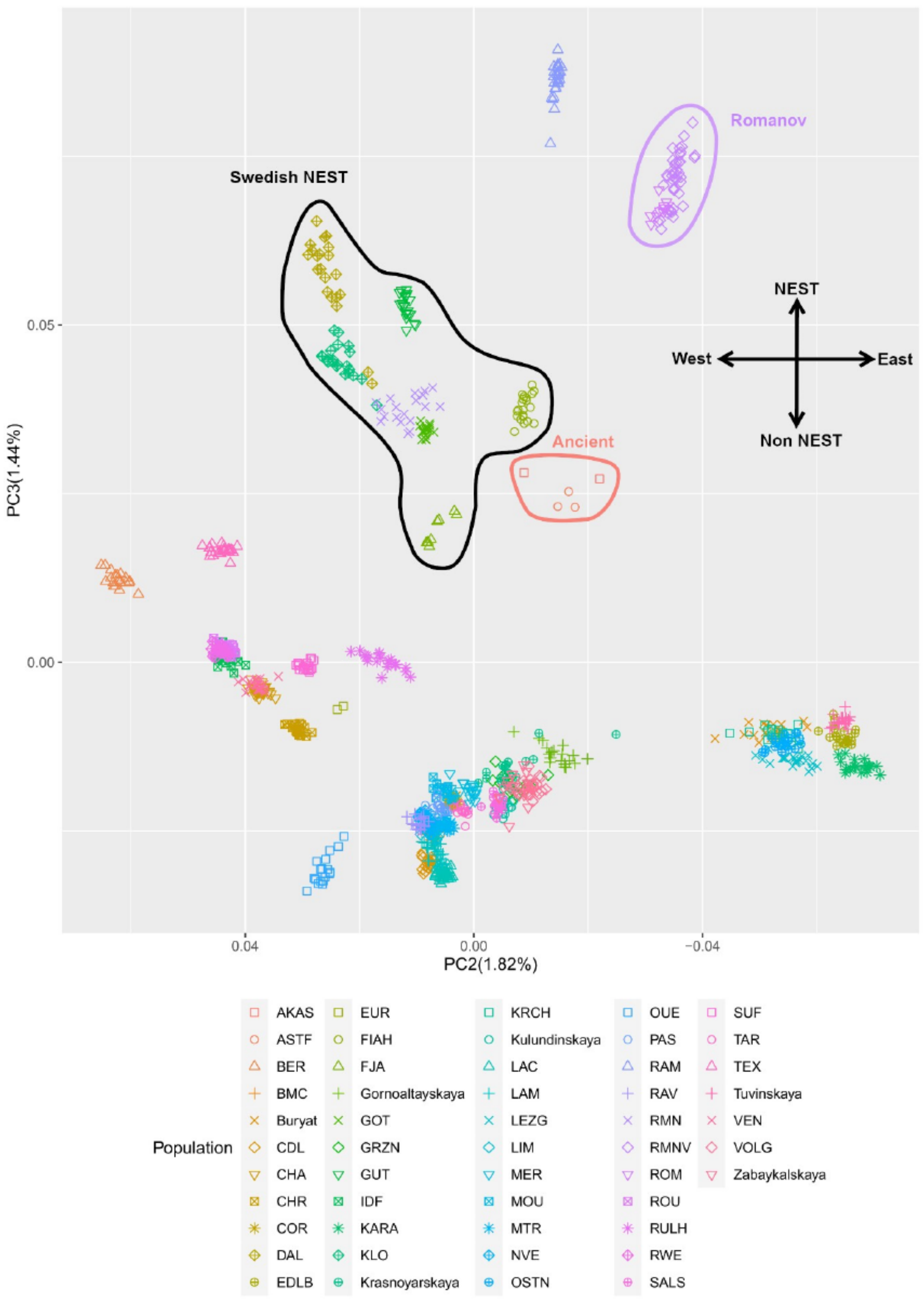


**Supplementary Figure 4:** SNPCHP PCA showing PC 2 vs PC 3, percentage of explained variation is shown within parentheses of axis titles. The Swedish NEST breeds, Romanov and the ancient samples are pointed out.

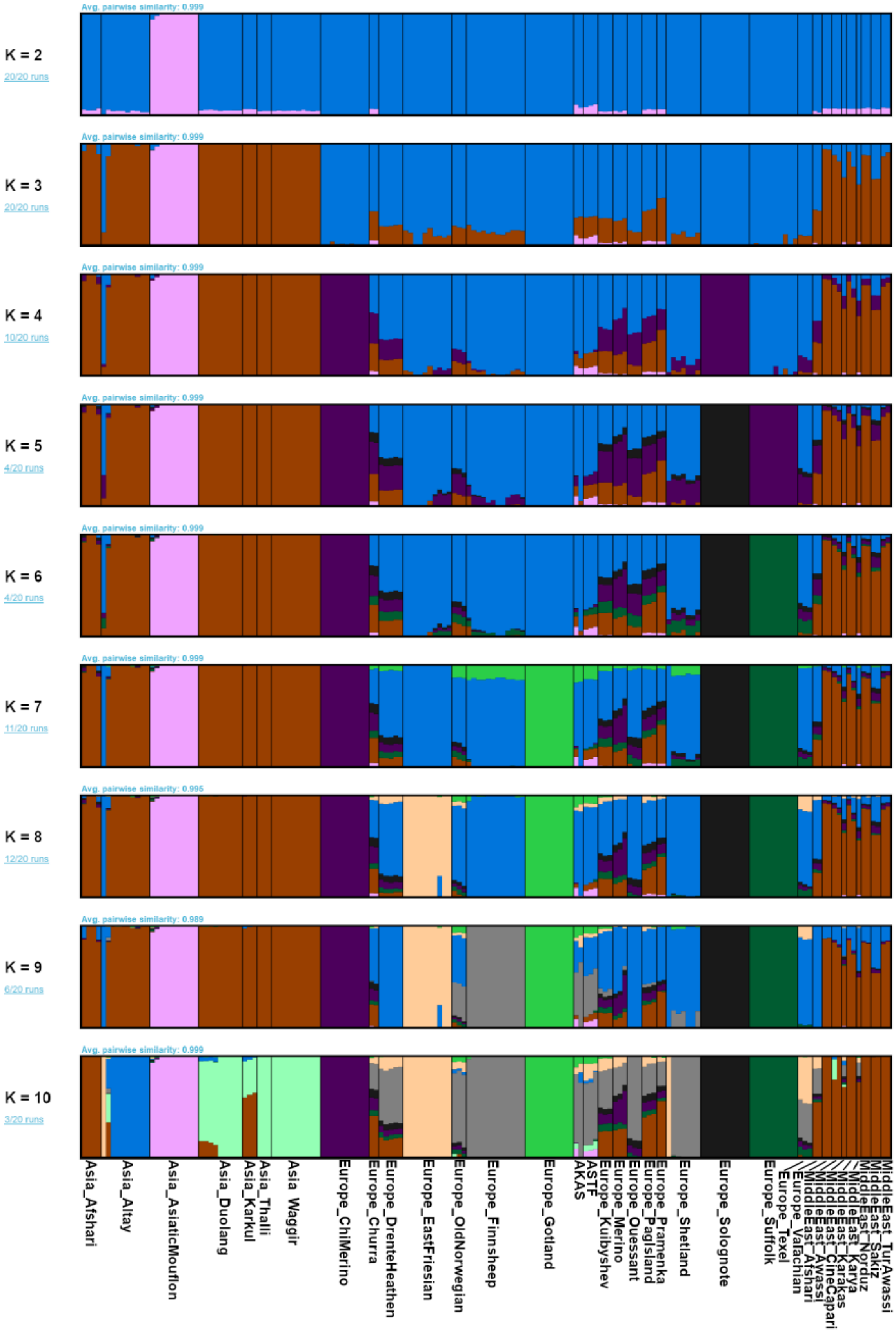


**Supplementary Figure 5**: WGS Results of 20 ADMIXTURE runs combined by PONG, Vertical bars indicate each individual's genetic variation belonging to a certain cluster, individuals are grouped by breed.

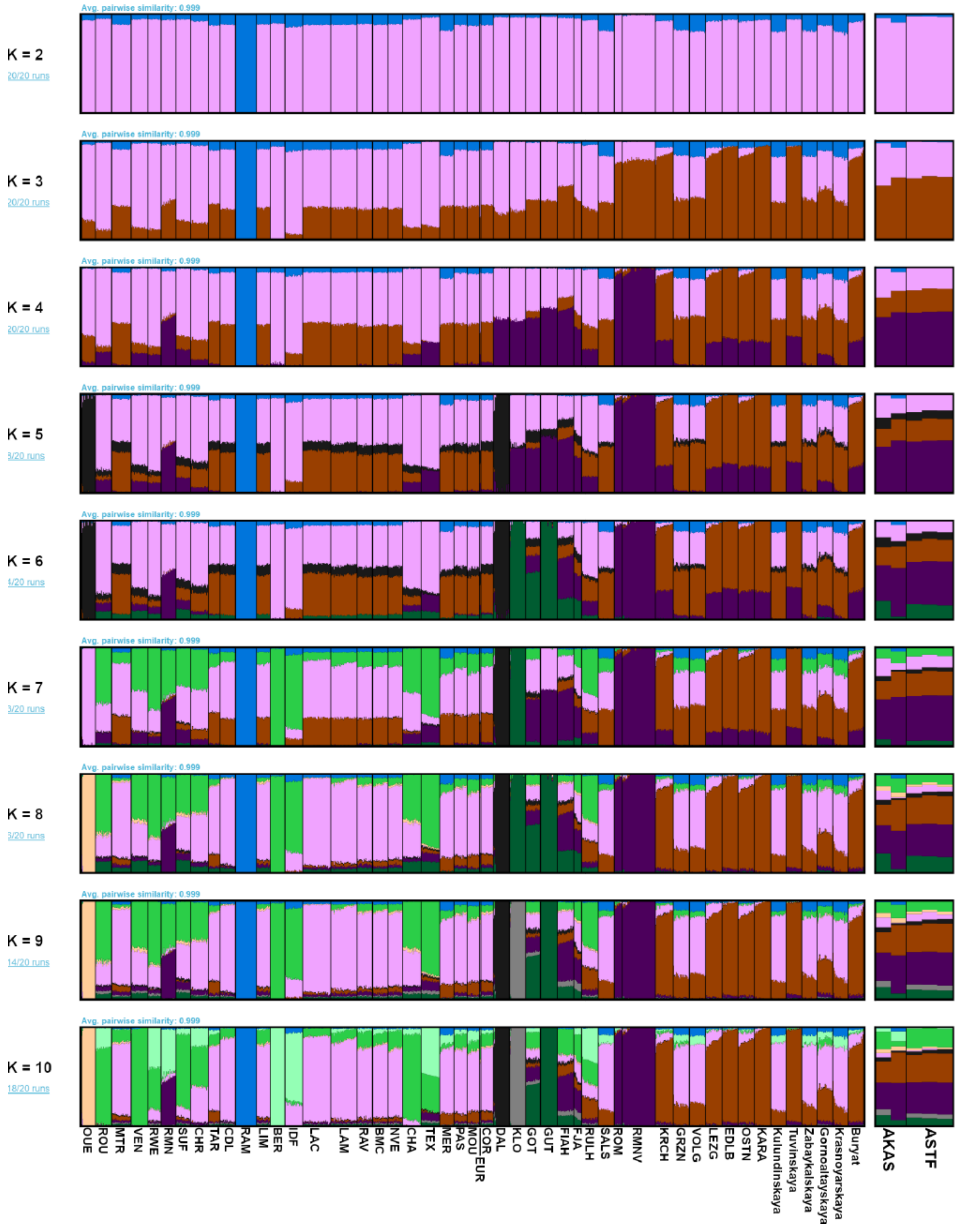

**Supplementary Figure 6**: SNPCHP Results of 20 ADMIXTURE runs combined by PONG, Vertical bars indicate each individual's genetic variation belonging to a certain cluster, individuals are grouped by breed.

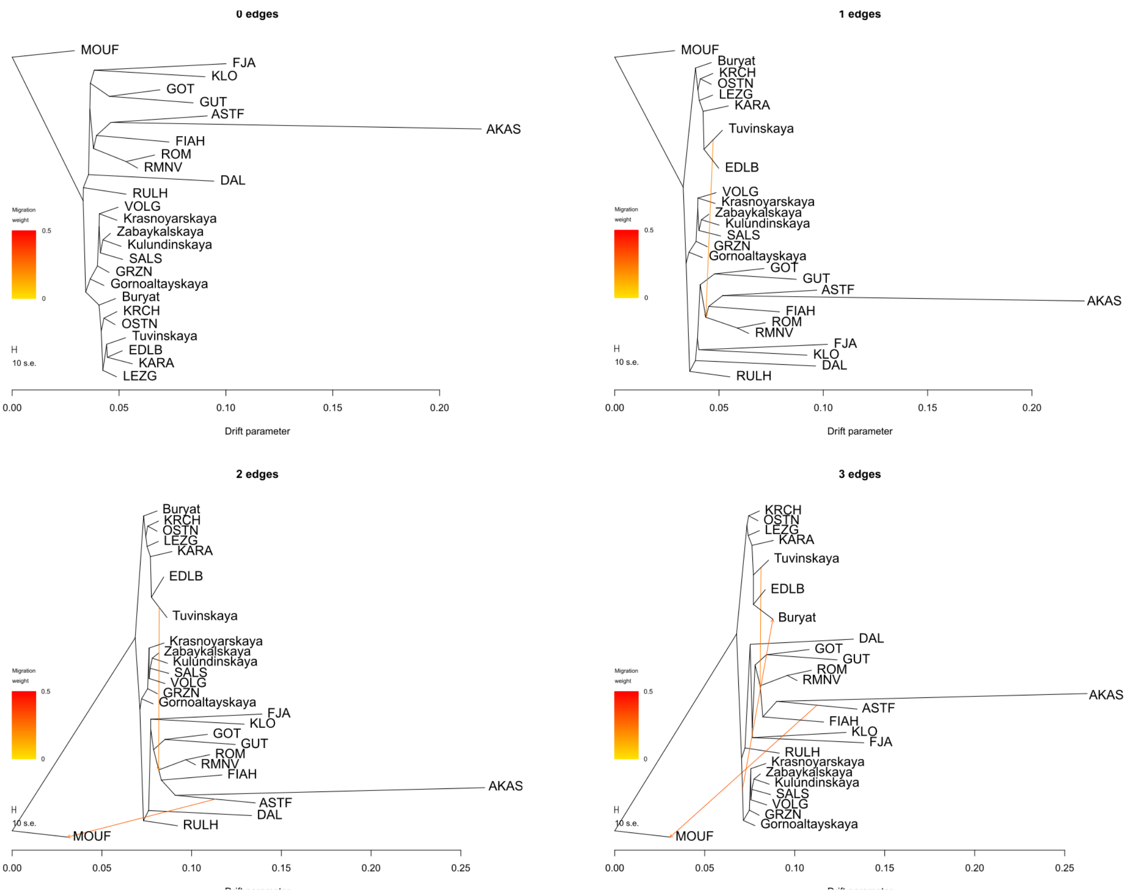


**Supplementary Figure 7**: OrientAGraph trees from the SNPCHP Panel, 0-3 migration edges, All Russian and Swedish breeds in SNPCHP panel, Asiatic mouflon as well as Ancients.

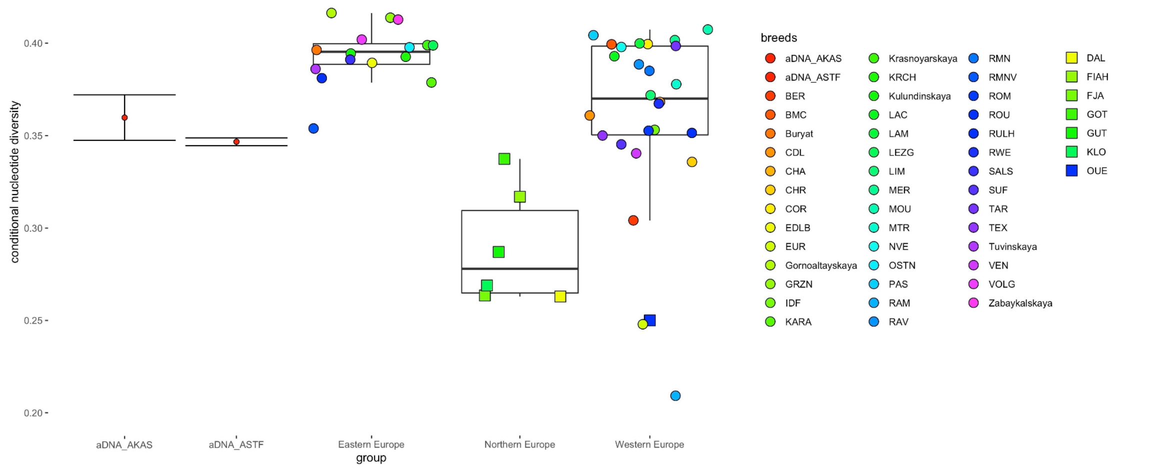


**Supplementary Figure 8**: Pairwise mismatch rate between randomly sampled individuals from the same group. This analysis was performed on the SNPCHP dataset.

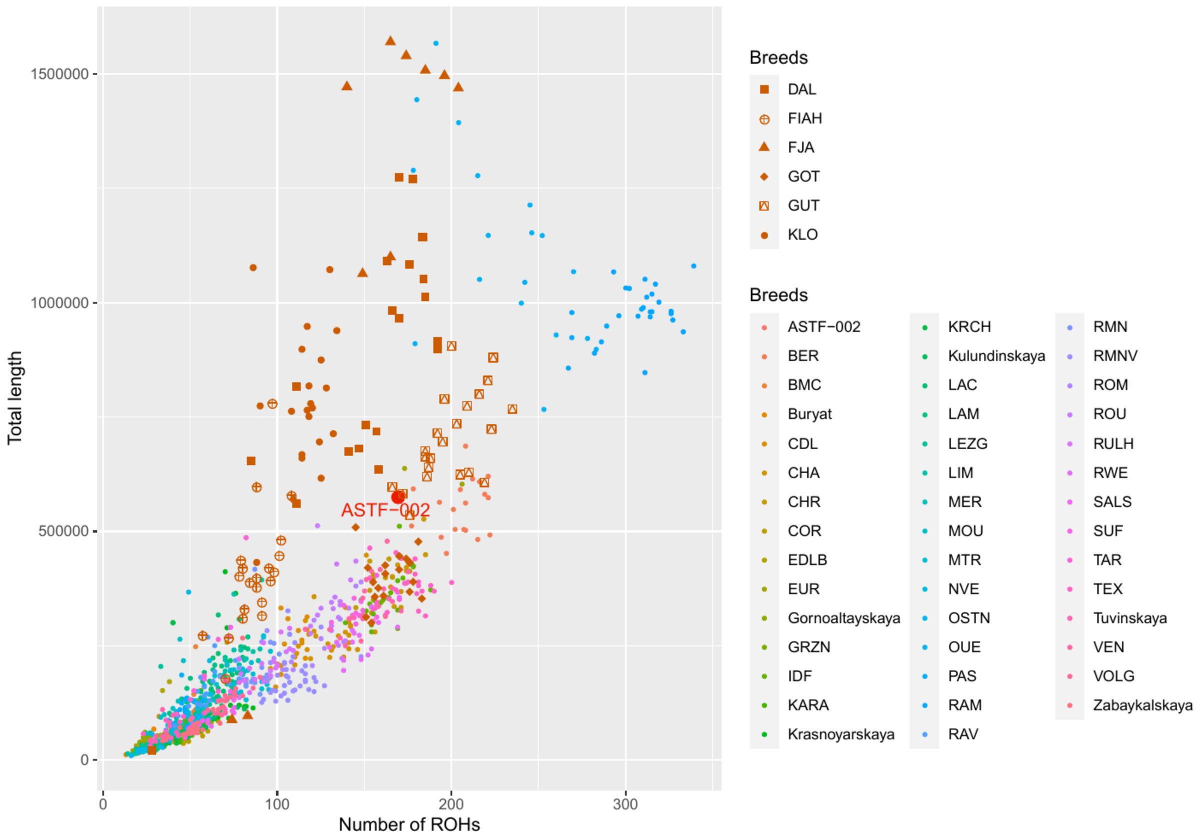


**Supplementary Figure 9**: Runs of homozygosity (ROH) for the SNPCHP dataset.

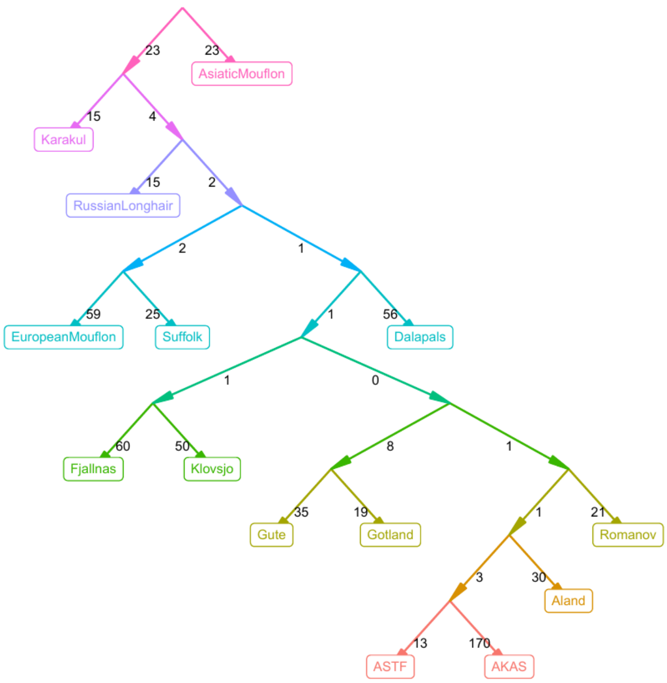


**Supplementary Figure 10a**: Best fitting admixture graph for m=0 and without constraints.

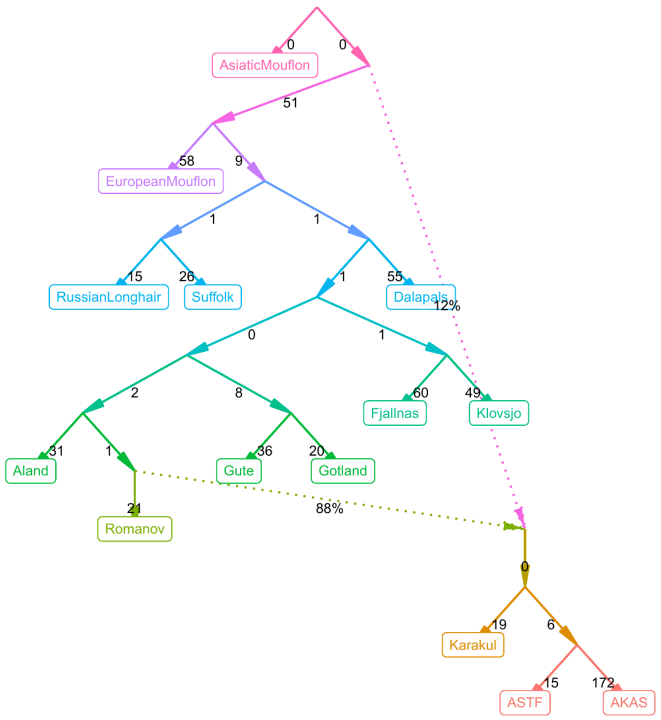


**Supplementary Figure 10b:** Best fitting admixture graph for m=1 and without constraints.

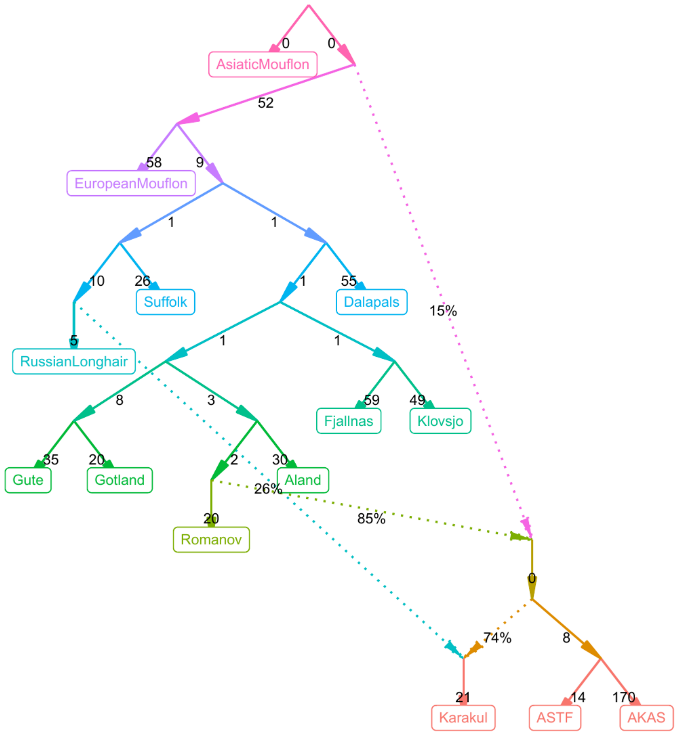


**Supplementary Figure 10c**: Best fitting admixture graph for m=2 and without constraints.

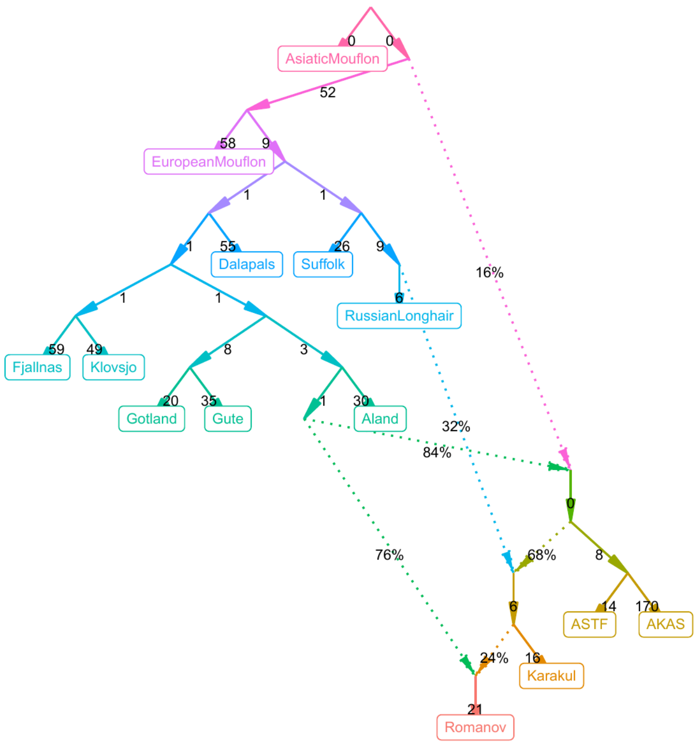


**Supplementary Figure 10d:** Best fitting admixture graph for m=3 and without constraints.

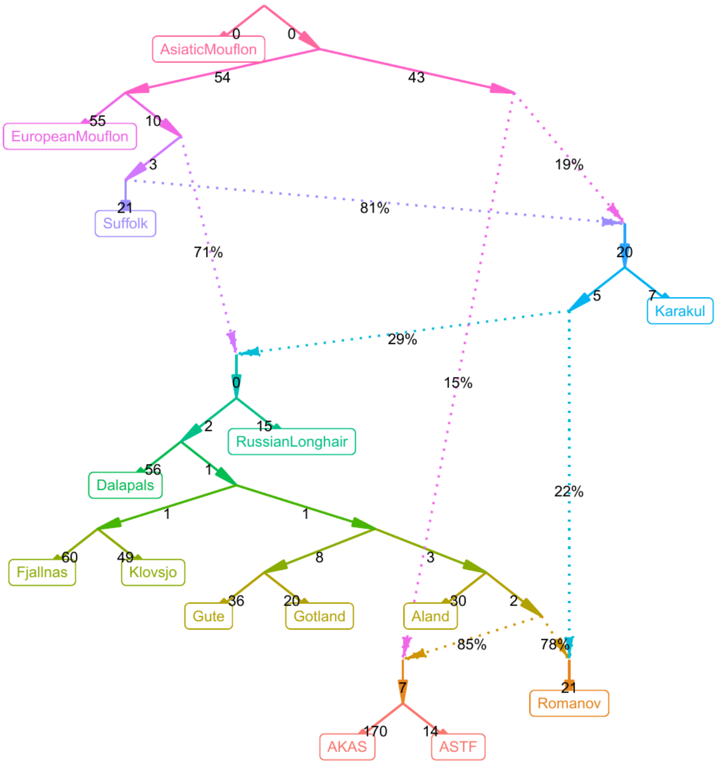


**Supplementary Figure 10e**: Best fitting admixture graph for m=4 and without constraints.

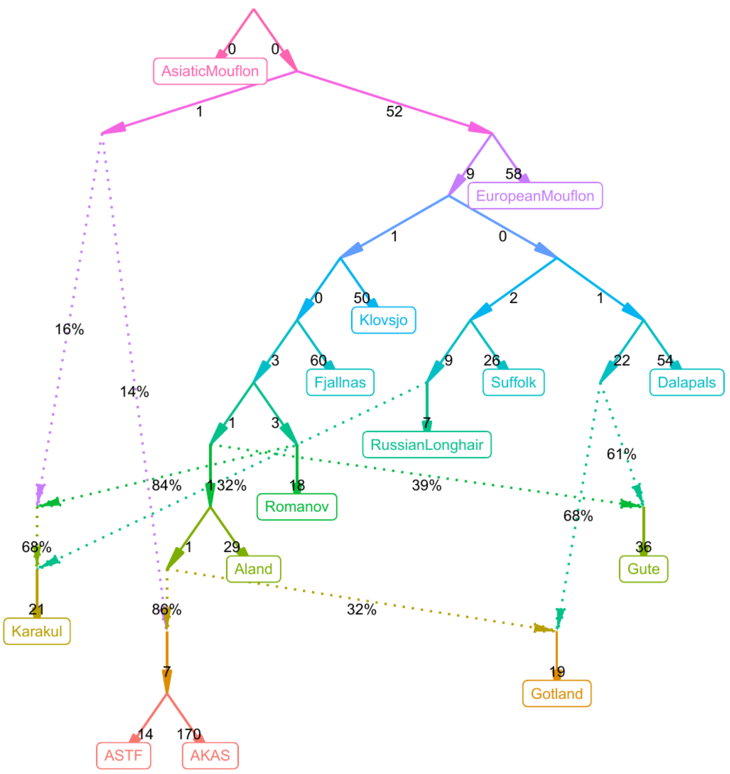


**Supplementary Figure 10f**: Best fitting admixture graph for m=5 and without constraints.

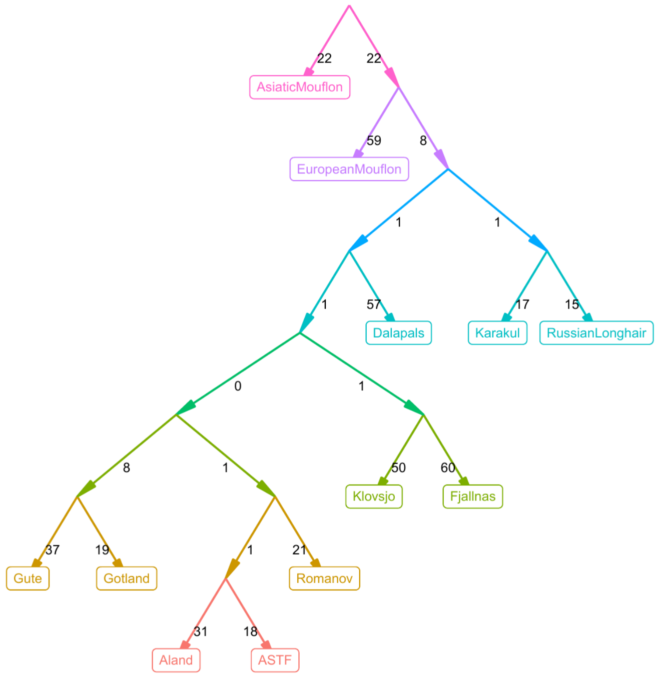


**Supplementary Figure 11a**: Best fitting admixture graph for m=0 and with constraints.

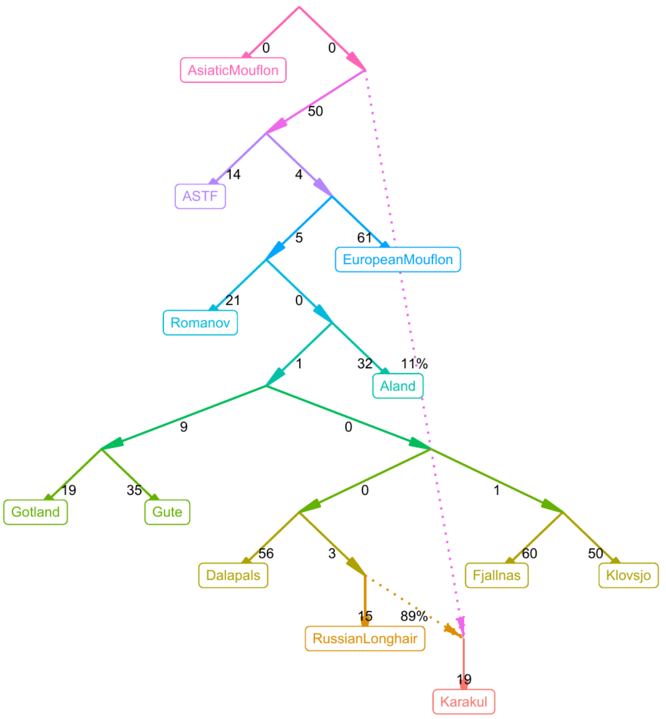


**Supplementary Figure 11b**: Best fitting admixture graph for m=1 and with constraints.

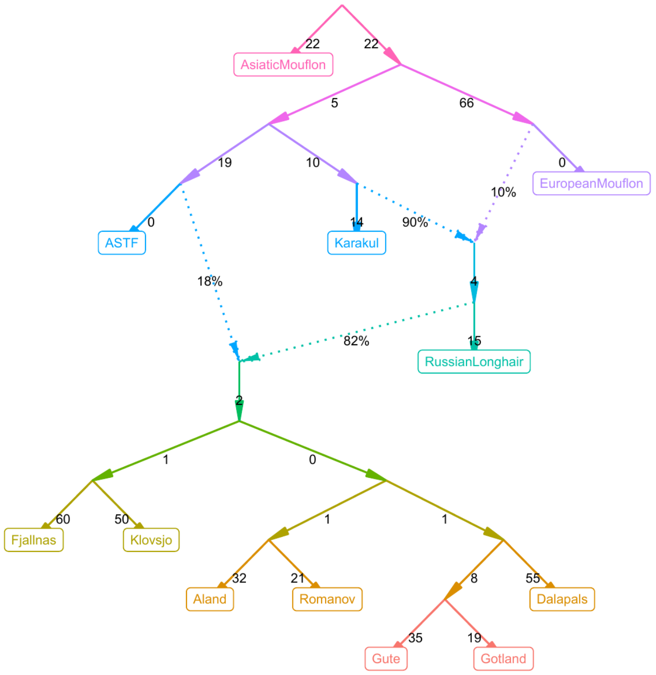


**Supplementary Figure 11c**: Best fitting admixture graph for m=2 and with constraints.

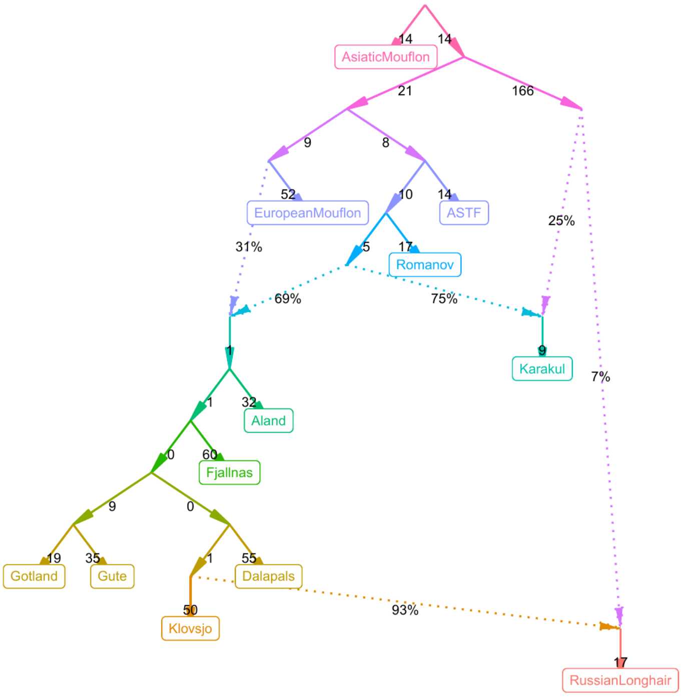


**Supplementary Figure 11d**: Best fitting admixture graph for m=3 and with constraints.

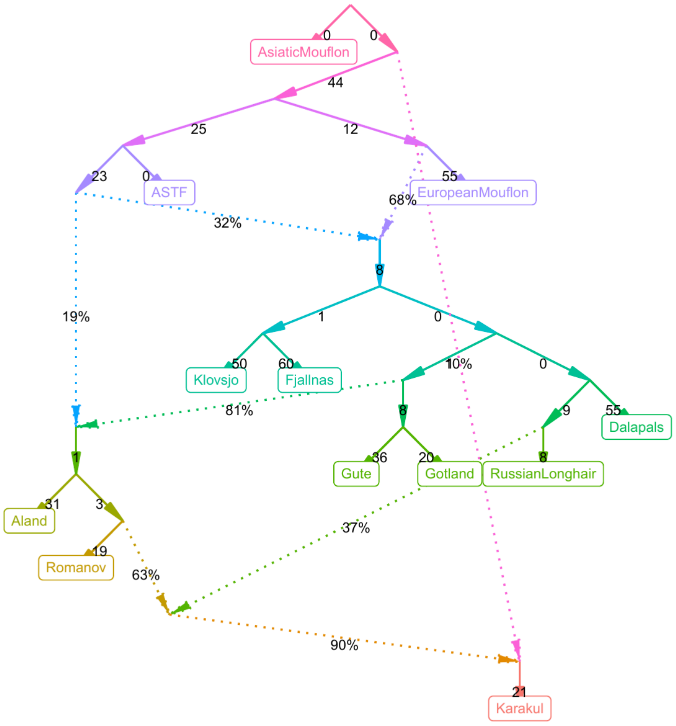


**Supplementary Figure 11e**: Best fitting admixture graph for m=4 and with constraints.

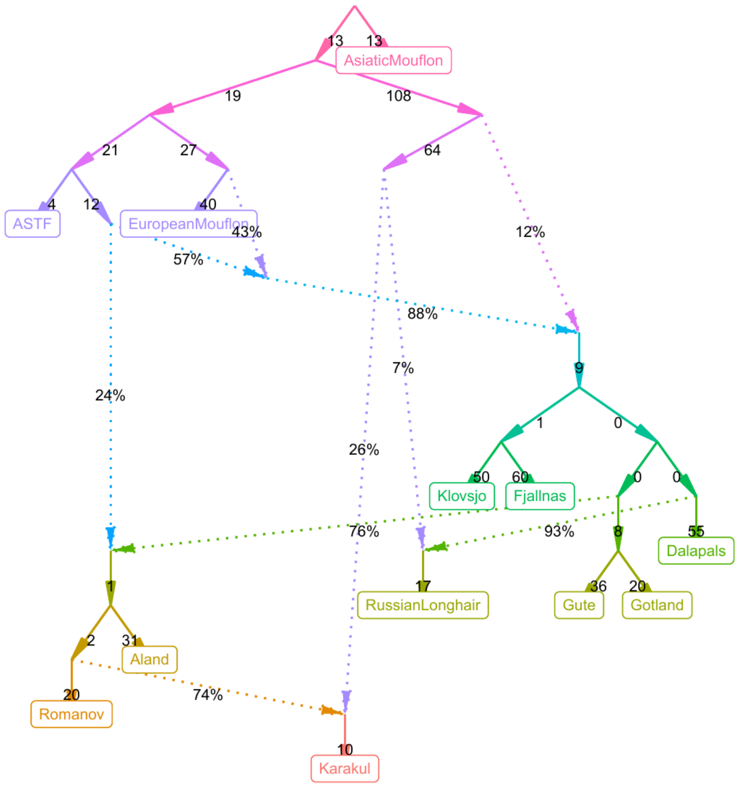


**Supplementary Figure 11f**: Best fitting admixture graph for m=5 and with constraints.

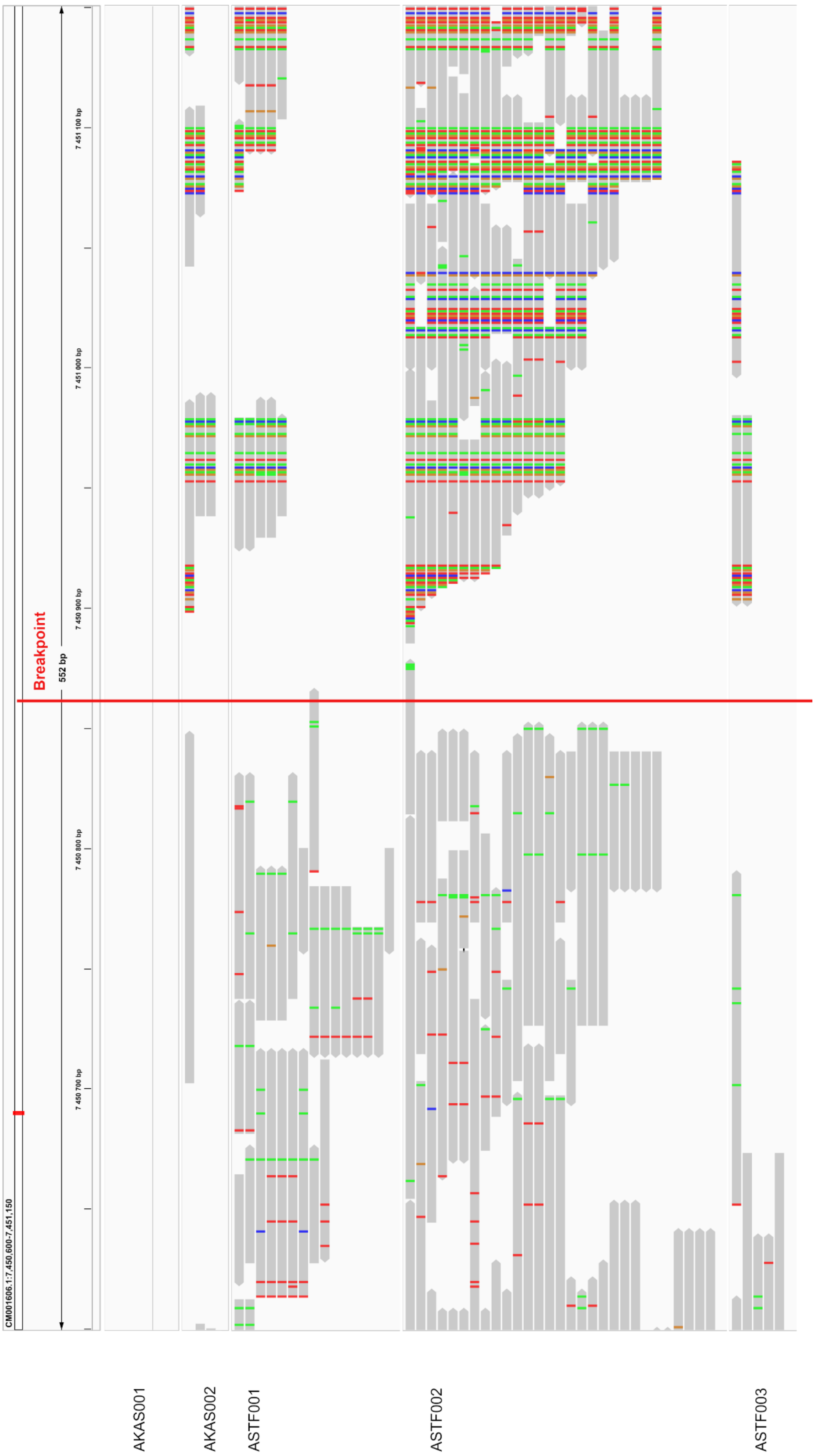


**Supplementary Figure 12a**: IGV (Robinson et al. 2011) plot of reads mapping to the ancestral (‘hairy’) reference genome version without the antisense *EIF2S2* retrogene into the 3’ untranslated region (UTR) of the gene *IRF2BP2*. The insertion point is marked by a red line.

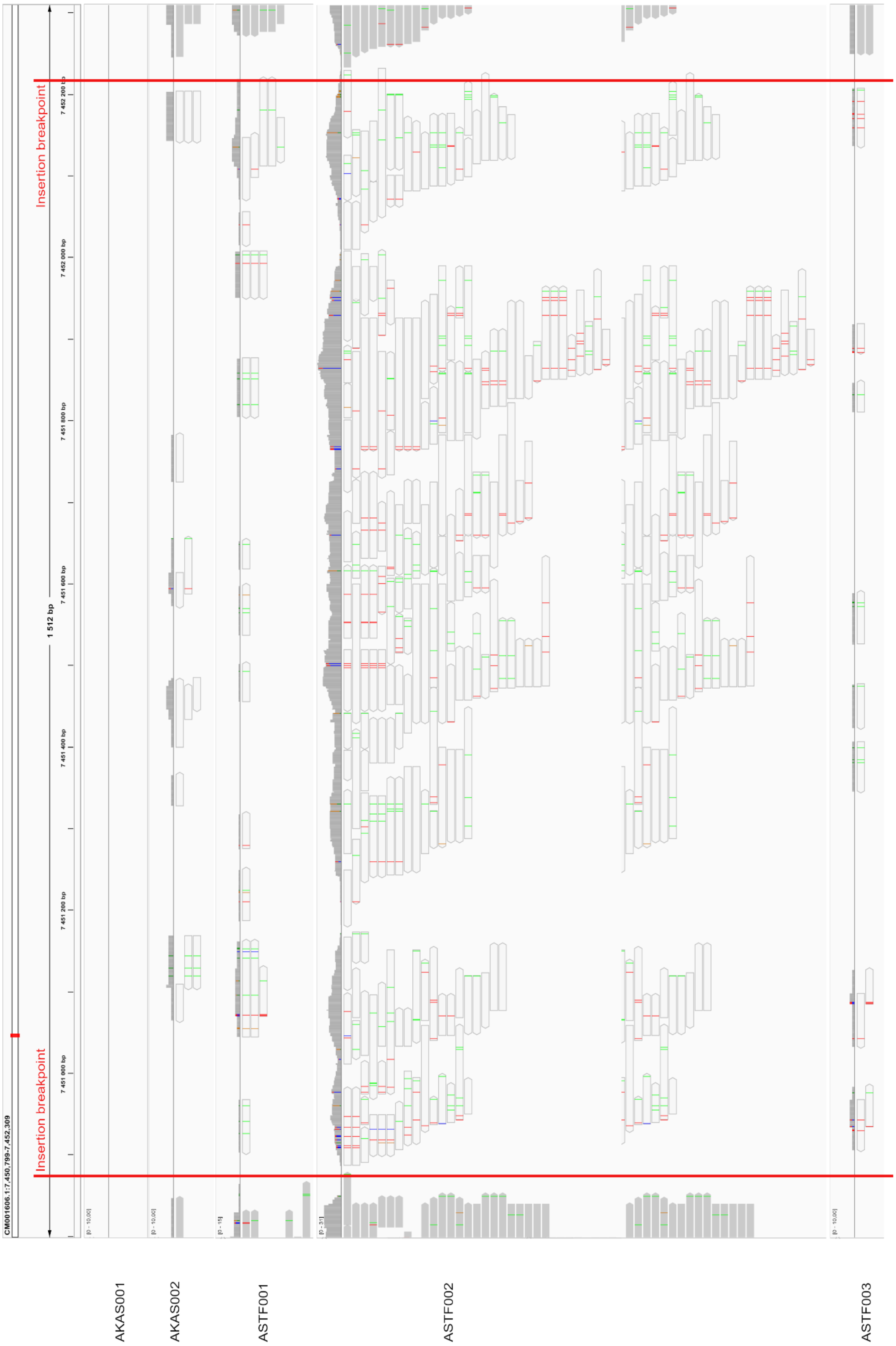


**Supplementary Figure 12b**: IGV plot (Robinson et al. 2011) of reads mapping to the derived ('wooly') reference genome version without the antisense *EIF2S2* retrogene into the 3' untranslated region (UTR) of the gene *IRF2BP2*. The insertion break points are marked by red lines. Reads shown in white have a mapping quality of 0, i.e. they map to multiple positions in the genome.

**Additional References in the Supplementary Information**


Popkin PRW, Baker P, Worley F, Payne S, Hammon A. 2012. The Sheep Project (1): determining skeletal growth, timing of epiphyseal fusion and morphometric variation in unimproved Shetland sheep of known age, sex, castration status and nutrition. *Journal of Archaeological Science* 39:1775–1792.

Robinson JT, Thorvaldsdóttir H, Winckler W, Guttman M, Lander ES, Getz G, Mesirov JP. 2011. Integrative genomics viewer. *Nature biotechnology* 29:24–26.
